# Monitoring Spore Dispersal and Early Infections of *Diplocarpon coronariae* Causing Apple Blotch Using Spore Traps and a New qPCR Method

**DOI:** 10.1101/2021.07.25.453640

**Authors:** Clémence Boutry, Anne Bohr, Sascha Buchleither, Mathias Ludwig, Thomas Oberhänsli, Lucius Tamm, Hans-Jakob Schärer, Pascale Flury

## Abstract

Apple blotch (AB) is a major disease of apple in Asia and recently emerged in Europe and the USA. It is caused by the fungus *Diplocarpon coronariae* (*Dc*) *(formerly: Marssonina coronaria;* teleomorph: *Diplocarpon mali*) and leads to severe defoliation of apple trees in late summer resulting in reduced yield and fruit quality. To develop effective disease management strategies, a sound knowledge of the pathogen’s biology is crucial. Data on the early phase of disease development is scarce: no data on spore dispersal in Europe is available. We developed a highly sensitive TaqMan qPCR method to quantify *Dc* conidia in spore trap samples. We monitored temporal and spatial dispersal of conidia of *Dc*, and progress of AB in spring and early summer in an extensively managed apple orchard in Switzerland in 2019 and 2020. Our results show that *Dc* overwinters in leaf litter and spore dispersal and primary infections occur in late April and early May. We provide the first results describing early-season dispersal of conidia of *Dc*, which, combined with the observed disease progress, helps to understand the disease dynamics and will be a basis for improved disease forecast models. Using the new qPCR method, we detected *Dc* in buds, on bark and fruit mummies, suggesting that several apple tissues may serve as overwintering habitats for the fungus, in addition to fallen leaves.

## INTRODUCTION

*Diplocarpon coronariae* (*Dc*) (Ellis & Davis) Wöhner & Rossman (Crous et al. 2020), formerly *Marssonina coronaria* (Ellis & Davis) Davis; teleomorph *Diplocarpon mali* (Harada & Sawamura) is an ascomycete fungus that causes apple blotch (AB) (Wöhner and Emeriewen 2019). The disease can result in severe tree defoliation and weakened trees, and ultimately decreased yield (Sharma and Thakur 2011) and fruit quality (Park et al. 2013). AB has a significant economic impact, especially in South and East Asia. In South Korea, the loss due to AB is estimated at US$ 29.79 M (Kwon et al. 2015). In India, AB is emerging as the most destructive disease affecting apple trees, becoming a major bottleneck in apple cultivation in Himachal Pradesh, an important apple producing state in the Western Himalayan region (Rather et al. 2017a; Sharma and Gupta 2018). Recently, AB has become an issue in Europe (Wöhner and Emeriewen 2019) and in the USA (Aćimović and Donahue 2018; Khodadadi and Aćimović 2019), especially in low-input orchards, for example organic orchards and untreated orchards used for juice production (Bohr et al. 2018; Hinrichs-Berger and Müller 2012; Persen et al. 2012). With the rising demand for reduced pesticide residues, the prevalence of AB may increase in conventionally managed apple orchards in the future.

To date, the biology and epidemiology of *Dc* and measures to control AB have been investigated primarily in Japan, China, Korea, and India, where the disease has been an issue since the late 1990s, while research on *Dc* and AB in Europe and the USA is nascent (Wöhner and Emeriewen 2019). *Dc* has a hemibiotrophic lifestyle (Horbach et al. 2011; Zhao et al. 2013) and the optimal conditions for infection are temperatures between 20 and 25°C and prolonged leaf wetness (Sastrahidayat and Nirwanto 2016). The minimum leaf wetness period for infection is eight hours at 15°C, and the risk of infection increases with increasing leaf wetness period and increasing temperature (Sharma et al. 2009). During the epidemic in summer the pathogen reproduces asexually by two-celled conidia. Additionally, in fall single celled microconidia (spermatia) are produced and can be found in acervuli together with conidia (Harada et al. 1974; Lindner 2012). Initial infections in spring can be caused by conidia released from acervuli or by ascospores developing in apothecia (Wöhner and Emeriewen 2019). However, the sexual stage of *Dc* has been described only in India, Japan, and China (Gao et al. 2011; Harada et al. 1974; Sharma and Gupta 2018), but not in Europe (Hinrichs-Berger 2015; Wöhner and Emeriewen 2019), the USA or Korea (Back and Jung 2014). The contribution of the different spore types to initial infections and their mode of dispersal has not been conclusively clarified, but wind as well as splash dispersal has been reported (Dong et al. 2015; Khodadadi et al. 2022; Kim et al. 2019). *Dc* overwinters in fallen leaves (Back and Jung 2014). Alternatively, buds and twigs have been hypothesised as overwintering refugia for *Dc* (Wöhner and Emeriewen 2019). However, data on the sources of primary infections and early disease progress is lacking in Europe and the USA.

To predict the risk of primary infections by *Dc* and subsequent outbreaks of AB, it is crucial to understand the spore dispersal pattern of the fungus. Fungal spores can be quantified using spore traps combined with subsequent quantification using visual (microscopic) or molecular methods. There are various types of spore traps, and depending on the mode of pathogen dispersal, the research question or the available resources, certain spore traps are better suited. Spore traps either collect spores passively by gravitational deposition or actively by sampling specific volumes of air and capturing the spores by impaction, impingement, filtration, virtual impaction, cyclone or electrostatic attraction (West and Kimber 2015). The spores are collected either onto a solid surface like agar on a Petri dish, filter paper, a double-sided adhesive tape, petroleum jelly (Vaseline)-coated tape, slides or rods, or electrostatic plastic film, or into Eppendorf tubes (Kim et al. 2018), or more rarely into a liquid. Spores captured by spore traps have traditionally been identified and quantified by microscopy (Andersen et al. 2009; Sterling et al. 1999). However, microscopy requires a spore trapping surface that can be examined under a microscope and a trained investigator to identify and count the specific fungal spores accurately. Newer methods combine microscopy with image recognition of spores and machine learning (Basso et al. 2020; Kilin et al. 2019). Furthermore, laser-based real-time optical particle counters (OPC) that detect particles in the air are being tested for real-time detection of spores in the air (Basso et al. 2020; Kilin et al. 2019). Besides optical methods, molecular methods are used to quantify fungal spores. Quantitative real-time polymerase chain reaction (qPCR), loop-mediated isothermal amplification (LAMP) (Notomi et al. 2015; Ren et al. 2021), serial analysis of gene expression (SAGE), and microarray technology (Aslam et al. 2017) are based on quantification of DNA, while the enzyme-linked immunosorbent assay (ELISA) (Kennedy et al. 2000), and the fluorescent antibody (FA) assay (Schneider et al. 2009) detect fungal proteins via antibodies. In recent years the qPCR method has become the most frequently used quantitative molecular method to detect and quantify fungal pathogens. It offers a more sensitive and specific quantification compared to microscopy, as it can detect low concentrations of a fungal pathogen against a background of particles and DNA from diverse organisms (Parker et al. 2014) and does not depend on visual identification of a fungus spore.

Information on spore dispersal along with observations on disease progress and combined with weather data can provide a basis to understand disease dynamics and develop disease forecast models. Forecast models provide a valuable tool for the efficient application of crop protection measures (Agrios 2005; Hardwick 1998). RIMPro (RIMpro, Amsterdam, The Netherlands) developed a first disease forecast model for AB in Europe based on literature information from Asia. Recently, *Dc* spore dispersal was investigated in Korea from June to October, employing spore traps and microscopy, and a disease forecast model was developed (Kim et al. 2019). However, the authors did not investigate spore dispersal in the spring, which is a period critical to understanding the primary infection and disease onset and which may be subject to efficacious disease management.

This study aimed to gain a better understanding of the early phase of AB disease in spring and summer. Thus, the specific objectives were to i) develop a qPCR method that allowed the quantification of *Dc* spores in spore trap samples, ii) investigate temporal and spatial aspects of *Dc* spore dispersal in relation to development of AB in the field in spring and early summer, and iii) identify potential sources of primary *Dc* inoculum.

## MATERIALS AND METHODS

### Spore traps used in this study

Conidia and ascospores of *Dc* have been reported to be dispersed by wind (Dong et al. 2015; Kim et al. 2019), while conidia are also reported to be splash-dispersed (Dong et al. 2015). Two trap types, the Mycotrap and the rotating-arm spore traps, were used to sample airborne spores in the field (Figs. 1D, E). Additionally, apple bait plants were used as in vivo spore traps (Fig. 1F). A detailed description of the spore traps is provided in Supplementary Materials and Methods S1. Briefly, the Mycotrap (Siegfried et al. 1996) (Fig. 1D) is an impaction spore trap and similar to the Burkard 7-day Recording Volumetric Spore Sampler, which was used to capture airborne conidia of *Dc* in Korea (Kim et al. 2019). The air is sucked in horizontally through a sampling orifice and impacts a trapping surface on a cylinder inside a chamber. The cylinder slowly rotates, completing one turn over seven days, allowing continuous sampling for the seven day period. The rotating-arm spore traps (Fig. 1E) were constructed based on the description of Quesada et al. (2018) (details in Supplementary Materials and Methods S2). The rotating-arm spore traps consisted of two rods connected to a battery-powered motor (spinart™ Lightweight Hanging Motor, 30 rpm). Each rod held a microscope slide on which was mounted a strip of Vaseline coated plastic film (Supplementary Materials and Methods S3) attached with 25 mm foldback clips (Maul, Bad König, Germany). The rotating-arm spore traps were covered with an aluminum shield as rain protection.

**Fig. 1.**
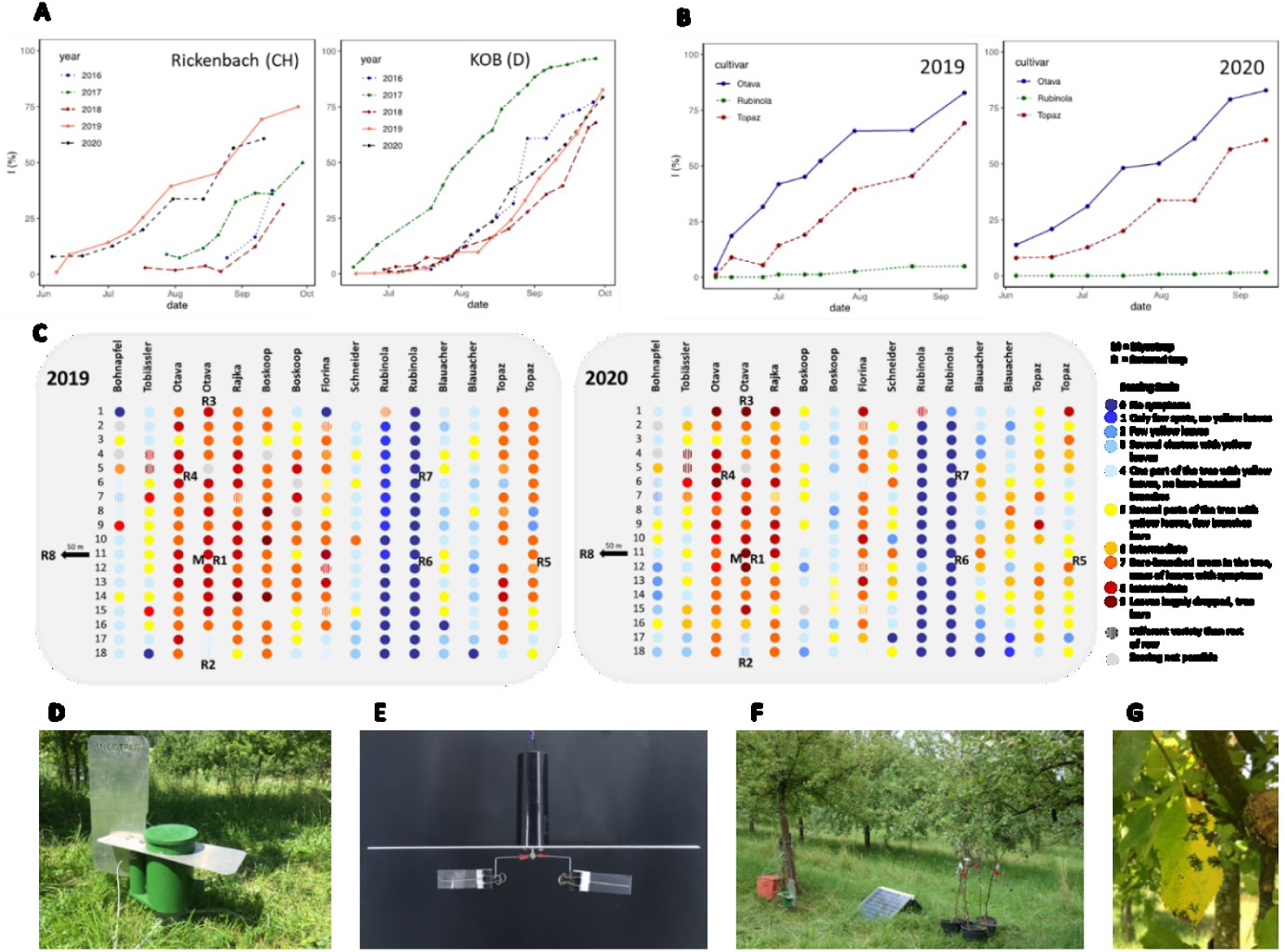
Apple blotch (AB) development in two apple orchards and the location of the spore traps. **A, B)** AB disease severity as percent damage according to the McKinney Index (I) (McKinney 1923). **A)** ‘Topaz’ trees in two orchards (Rickenbach, Switzerland and KOB, Germany) were regularly scored for severity of AB from June to October for five consecutive years (Rickenbach, 36 trees; KOB, 120 trees). **B)** AB development on ‘Topaz’ (36 trees), ‘Rubinola’ (35 trees), and ‘Otava’ (33 trees) in the Rickenbach orchard in 2019 and 2020. **C)** AB disease scoring in the Rickenbach orchard by middle of September 2019 and 2020. The orchard is planted with tree rows of ten different cultivars. Each dot represents one tree. The color of the dot indicates the disease score of the tree. The placement of the Mycotrap and rotating-arm spore traps is indicated by the letters M and R, respectively. One rotating-arm trap was installed 50 m outside the orchard (R8). **D, E, F)** Pictures of the spore traps used in this study. **D)** the Mycotrap, **E)** the rotating-arm spore trap, **F)** potted bait plants. **G)** leaf with typical AB symptoms.

To collect splash-dispersed *Dc* conidia in rain water, funnels were placed in Schott-bottles to collect rain splash. Approximately 100 mL of rainwater was collected per Schott-bottle and filtered through a cellulose acetate filter (25 mm in diameter, 0.8 μm pore size, Sartorius-Stedim, Goettingen, Germany) in polycarbonate filter housings. The filters were subsequently cut into pieces of approximately 3 × 3 mm and stored at -20°C.

### Design and evaluation of qPCR primers and probes

We aimed at developing a highly specific and sensitive TaqMan® qPCR assay for *Dc*. The primers and hydrolysis probes designed targeted the nuclear ribosomal internal transcribed spacer (ITS1) region between the 18S and 5.8S rDNA of *Dc* using the program Beacon Designer (V8.16, Premier Biosoft, Palo Alto, CA, USA) with the following parameters: amplicon length between 100 and 200 bp, melting temperature (TM) of primers at 60.0°C +/- 1.0°C, and TM of the probe at 10.0°C +/- 5.0°C above TM of the primers (Table 1). The program’s default settings were used for maximum ΔG of self-complementarity, 3’-end stability, and % GC. The specificity of the primers and probes was confirmed in silico using the National Center for Biotechnology Information (NCBI, Bethesda, MD, USA) Primer-Blast (Ye et al. 2012) by alignment of different ITS gene bank accessions of *Dc*.

**TABLE 1.**
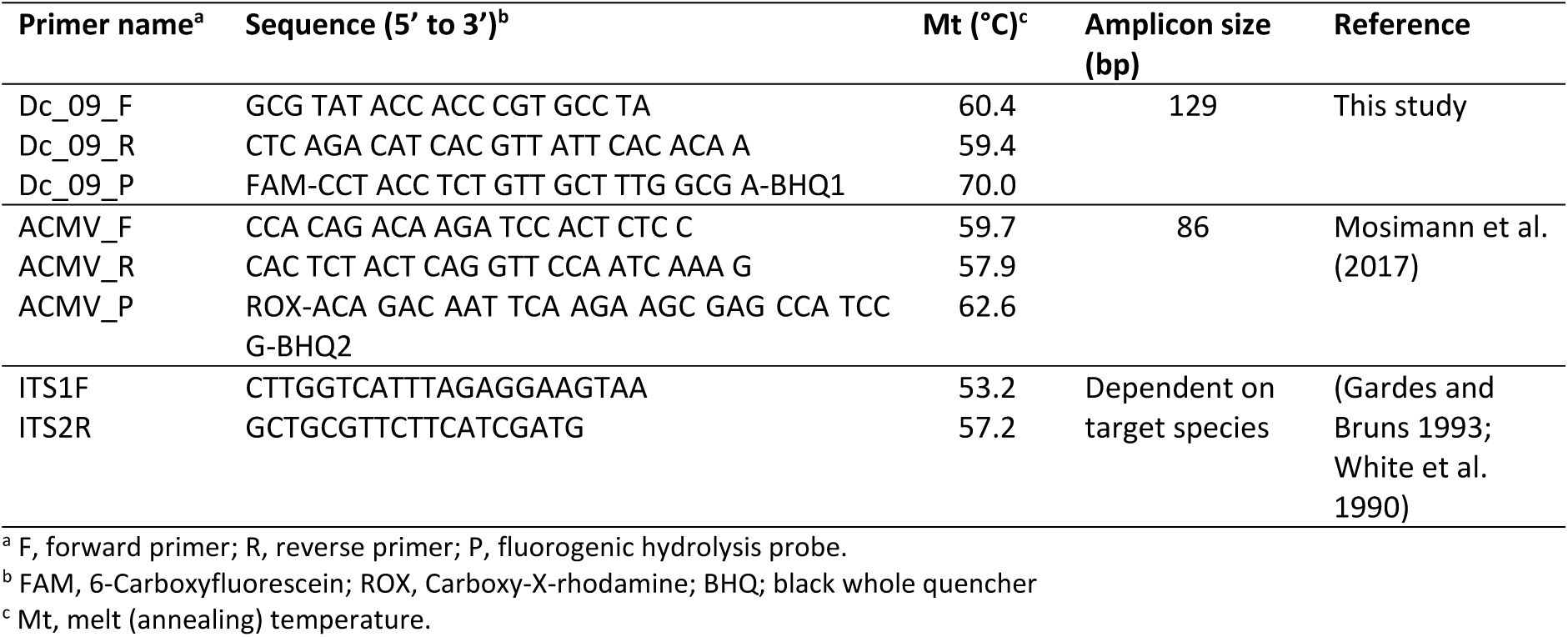
Primers used in the qPCR to detect DNA of *Diplocarpon coronariae*.

The specificity was further tested in vitro with genomic DNA of *Dc* isolates from Switzerland, Korea and Japan, with apple DNA, and with DNA of different fungi, i.e. fungi within the genus *Diplocarpon*, fungi causing Marssonina diseases on other hosts than apple, and fungi that are common pathogens of apple. Information on the tested isolates is provided in Supplementary Table S1. *Dc* isolates were cultured on potato peptone carrot dextrose agar (PPCDA) adapted from Zhao et al. (2010) and containing 100 mL carrot juice, 39 g potato dextrose agar, 10 g peptone, and 1 L double-distilled water. Other pathogens were cultivated on potato dextrose agar (PDA). Fungal DNA from the cultures was extracted using a ZymoBIOMICS® Quick-DNA Fungal/Bacterial Miniprep Kit (Zymo Research, Irvine, CA, USA) according to the manufacturer’s recommendations.

The qPCR was designed for use with a TaqMan® probe, but we also tested whether the primer pair Dc_09 could be used with a DNA intercalating fluorophore (SYBR Green).

### qPCR reaction conditions

All primers and probes were synthesized and purified by high-performance liquid chromatography (HPLC) at Microsynth AG (Balgach, Switzerland). Primers and probes (Table 1) were dissolved in a TE-dilution buffer (TE-Buffer: 10^−3^ mol ·L^−1^ Tris, 10^−5^ mol ·L^−1^ Na_2_EDTA, pH 8.0).

TaqMan® based qPCR reactions consisted of 5 μL KAPA PROBE FAST (Sigma-Aldrich Chemie AG, Buchs, Switzerland), 1 μL of each primermix for *Dc* and APA9 (containing primers at a concentration of 3 μM for Dc_09 and 2 μM Africa Cassava Mosaic Virus (ACMV), and the probe at 1 μM each), 0 to 2 μL double-distilled water and 1 to 3 μL of template DNA in a total volume of 10 μL. SYBR based *Dc* specific qPCR reactions consisted of 5 μL KAPA SYBR FAST (Sigma-Aldrich Chemie AG, Buchs, Switzerland), 1 μL primermix (forward and reverse primers at a concentration of 3 μM each), 3 μL double-distilled water and 1 μL of template DNA in a total volume of 10 μL. The qPCR assay and data analysis were performed using a CFX96 Touch Real-Time PCR Detection System (Biorad, Hercules, CA, USA). The amplification and quantification conditions used were an initial denaturation step of 3 min at 95°C, followed by 39 and 45 cycles of 10 s at 95°C, and 20 s at 60°C for SYBR and TaqMan® qPCR, respectively. After each SYBR qPCR, a dissociation curve analysis was performed by gradually increasing the temperature from 65°C to 95°C by 0.5°C per cycle.

In specificity tests, a SYBR green qPCR assay was performed with the ITS primers ITS1F/2R in addition to the *Dc* specific qPCR, which served as an amplification control detecting all tested fungal species (Table 1). The reactions consisted of 5 μL KAPA SYBR FAST, 1 μL primermix (primers at a concentration of 2 μM each), 3 μL double-distilled water and 1 μL of template DNA in a total volume of 10 μL and were subjected to an initial denaturation step of 10 min at 95°C, followed by 39 cycles of 30 s at 95°, 30 s at 50°C, and a final extension step for 1 min at 72°C.

To visualize the PCR products, 10 μL of the qPCR reaction were loaded on a 2% Tris-acetate-EDTA buffered agarose gel stained with ROTI® GelStainRed (Carl Roth GmbH, Karlsruhe, Germany) and run at 50 V for 60 min. Images were captured under u.v. using an Azure Biosystems c150 (Azure Biosystems, Dublin, CA, USA) gel imaging workstation.

### DNA extraction from spore trap samples and apple leaves

To extract DNA from *Dc* conidia from Vaseline coated plastic film (Mycotrap and rotating-arm trap samples, Supplementary Materials and Methods S3) we used the ZymoBIOMICS® Quick-DNA Fungal/Bacterial Microprep Kit (Zymo Research, Irvine, USA) following the manufacturer’s protocol with two modifications: the Bashing Bead® tubes from the Zymo Microprep Kit were replaced by 2 mL tubes with screw caps containing 100 mg of 0.5 mm Zirconia bashing beads (N034.1 Roth, Karlsruhe, DE) and the Bashing Bead™ Buffer volume was reduced from 750 μL to 550 μL.

DNA from filter pieces (rain-splash samples) was extracted using a ZymoBIOMICS® Quick-DNA Fungal/Bacterial Microprep Kit following the manufacturer’s protocol, but adding skim milk to the bashing bead buffer at a final concentration of 2% to prevent the adhesion of free DNA to the filter membrane (Liang and Keeley 2013).

DNA from apple leaves with ambiguous AB symptoms was extracted following a CTAB buffer protocol (described in Supplementary Materials and Methods S4).

### qPCR based quantification of *D. coronariae* spores with a standard curve

The number of conidia corresponding to a given C_q_ value was calculated using a standard curve based on known quantities of conidia. Conidia suspensions were prepared by stirring AB-symptomatic apple leaves with sporulating *Dc* for 10 min in Volvic® natural spring water (Danone S.A. Paris, France), which is routinely used in our lab for the preparation of spore suspensions, and subsequently filtering the spore suspension through a sieve (mesh size 0.1 mm). The spore concentration was assessed using a haemocytometer (0.1 mm depth, 0.0025 mm^2^, Neubauer®, Paul Marienfeld GmbH, Lauda-Königshofen, Germany). A ten-fold dilution series was prepared in double-distilled water containing 0.05% Tween® 80 as described by Mc Devitt et al. (2004), and 10^5^ to 10^1^ conidia were pipetted to the material used in spore traps (Vaseline coated plastic film or filter pieces) and let dry. DNA extraction and qPCR were performed as described above. The standard curves were used for the test calibration, the limit of quantification (LOQ), and the calculation of spore numbers based on Cq values.

All spore trap samples were spiked with 10^7^ copies of linearized APA9 plasmid (vector pUC19 with an insert ACMV; NCBI GenBank accession number AJ427910) (details in the Supplementary Materials and Methods S5) (Mosimann et al. 2017) as an internal control to account for differences in DNA extraction efficiency between different samples. The APA9 plasmid was quantified together with *Dc* using a multiplexed TaqMan® qPCR with the primers and probe for ACMV listed in Table 1. For multiplexing, the singleplex TaqMan® qPCR protocol for *Dc* described above was adapted by replacing 1 μL of double-distilled water with 1 μL primermix for APA9 (containing primers at a concentration of 2 μM and the probe at 1 μM). The results were used to normalize DNA extraction using the method of Von Felten et al. (2010).

### Field sites

*Dc* spore dispersal and AB epidemiology were investigated in an extensively managed organic apple cider orchard in Rickenbach (Zurich, Switzerland, 47°33’30.5”N 8°47’29.9”E). The site is characterized by a mean annual temperature of 10.0°C and mean annual rainfall of 980 mm (years 2010 to 2020). The orchard was planted in 2002 to ten different apple cultivars arranged in 15 tree rows with 18 apple trees in each row (Fig. 1C) and an in-row planting distance of 4 m and between-row planting distance of 9 m. In 2019, the orchard was treated with organic fungicides (acidified clay minerals, Myco-Sin® at 8 kg/ha, Andermatt Biocontrol Suisse AG, 6146 Grossdietwil, Switzerland; in a tank mix with Sulphur, Netzschwefel Stulln at 4 kg/ha, Biofa AG, Münsingen, Germany) on 25 May. The cultivars ‘Otava’, ‘Rajka’, ‘Rubinola’, ‘Topaz’, and ‘Boskoop’ received additional sulfur with lime (Curatio®, Biofa AG, Münsingen, Germany) on 11 June. In 2020, the orchard was treated with sulfur and clay on 27 April, 9 May, 22 May (all cultivars except ‘Bohnapfel’, ‘Schneider’, and ‘Tobiässler’), and on 2 June (only ‘Otava’, ‘Rajika’, ‘Rubinola’, and ‘Boskoop’). Trees with spore traps were excluded from receiving fungicide treatments in 2019 and 2020. In addition, in 2020, the first tree of each row was left untreated.

AB epidemiology was further investigated in an organic apple orchard at the Competence Center for Fruit Crops at Lake Constance (Kompetenzzentrum Obstbau Bodensee (KOB), Ravensburg, Germany, 47°46’06.8”N 9°33’18.0”E). The site is characterized by a mean annual temperature of 9.7°C and mean annual rainfall of 908 mm (years 2010-2020, data from the weather station in Bavendorf, www.wetter-bw.de). The orchard was planted in 2003 to ‘Topaz’ on M9 rootstocks with a 3.2 m between-row and 0.8 m in-row planting distance. For monitoring AB, 120 trees in two consecutive rows were regularly assessed. No fungicide treatments were applied to the trees assessed during the entire period of the study.

Weather data including rainfall, leaf wetness, and temperature were obtained from weather stations close to the field sites: the Agrometeo (Agroscope, Nyon, Switzerland; www.agrometeo.ch) weather station “Liebensberg”, Zurich, Switzerland (47°53’38.1”N 8°83’73.8”E, altitude 535 m), and the Agrarmeteorologie Baden-Württemberg (Landwirtschaftliches Technologiezentrum Augustenberg, Tübingen, Baden-Württemberg, Germany, www.wetter-bw.de) weather station “Bavendorf”, Germany (47°76’84.4”N 9°55’99.0”E, altitude 481 m).

### Disease assessment in the apple orchard

From 2016 to 2020, the severity of AB was assessed every one to two weeks from the onset of first symptoms through October by scoring entire trees using a qualitative ordinal scale of 0 to 9 (0 = no disease symptoms; 1, only few spots, no yellow leaves; 2, few yellow leaves; 3, several clusters with yellow leaves; 4, one part of the tree with yellow leaves, no bare branches; 5, several parts of the tree with yellow leaves, few branches bare; 6, intermediate; 7, bare-branched areas in the tree, mass of leaves with symptoms; 8 intermediate; 9, leaves largely dropped, tree bare).

The ordinal scores were used to calculate damage (in percent) based on the McKinney Infection Index (I) (McKinney 1923), where I (%) = [sum (class frequency × score of rating class)]/[(total number of ratings) × (maximum grade)] × 100. The scoring scale and the results for 2016 to 2018 have been presented previously in non-peer-reviewed journals for practitioners (Schärer et al. 2019; Wöhner et al. 2019).

### Monitoring spore dispersal in an apple orchard

To investigate dispersal of *Dc* conidia in the field, different spore traps were installed in the Rickenbach orchard from May to July and from March to July in 2019 and 2020, respectively. Exact sampling periods for each trap are provided in Supplementary Tables S2 and S3. To monitor the temporal resolution of the spore dispersal, one Mycotrap was placed on the ground in the previous year’s AB hotspot in the orchard within a row of ‘Otava’ trees (Figs. 1C, D). A second Mycotrap was placed in the canopy of an ‘Otava’ tree (2.5 m above ground) above the first Mycotrap. We hypothesized that primary spores originating from leaf litter on the ground might be detected rather by the trap on the ground, while a trap in the tree crown would catch first spores released from other sources. We also hypothesized that later in the season, when the leaves are heavily infected, there would be more spores in the crown than on the ground. Container-grown, two-year-old apple trees were used as bait plants and were placed for periods of five to fourteen days in the orchard and subsequently incubated under rain protection to assess disease development and correlate the presence of spores in the orchard with actual infections (Fig. 1F). In 2019, each series of bait plants consisted of three ‘Topaz’, three ‘Gala’ and three ‘Kiku’ apple trees. In 2020, each series of bait plants consisted of five ‘Topaz’ apple trees.

To observe the spatial gradients of airborne spore catches in relation to disease severity in the orchard, seven rotating-arm spore traps were mounted inside and at the edge of the orchard on trees with severe (3 ‘Otava’, 1 ‘Topaz’ tree), mild (1 ‘Schneider’ tree) and no *Dc* symptoms (2 ‘Rubinola’ trees) at one of the lowest branches of the apple trees (1.7 m above ground) and close to the trunk. Moreover, one rotating-arm spore trap was installed at the same height but approximately 50 m outside the orchard perimeter. The exact location of the traps is indicated in Fig. 1C. In 2019, the traps were installed only between 22 Mai and 5 June. However, in 2020, we covered the entire sampling period from March to July. The spore trap samples were analyzed by the TaqMan® qPCR method described above.

To observe the spatial gradients of splash-dispersed *Dc* conidia 12 Schott-bottles with a funnel were placed within the row directly below trees as well as between the rows under the open sky and outside the orchard. The spore trap positions differed regarding disease severity of nearby trees. Rainwater was collected for 16 hours from 24 to 25 September 2020. It was the first rainy period after seven consecutive days without precipitation. The rainfall registered for the 16-hour period was 14 mm.

### Precipitation and spore number

To assess whether spore number and precipitation are correlated, a Kendall rank correlation test was performed in RStudio version 2021.09.1+372 (http://www.rstudio.com) using R version 4.1.2. with the spore number as dependent variable and the precipitation as independent variable. Since the origin and amount of the inoculum changed over time the analysis was performed for each month separately.

### Investigating possible sources of primary inoculum

To understand *Dc* spore dispersal, it is essential to know all sources of primary inoculum. We first studied spore dispersal from a deposit of overwintered *Dc* infected leaf litter. In 2019, a Mycotrap was placed in the field on a mesh cage depot filled with heavily *Dc* infested leaf litter of the variety ‘Remo’ at the Competence Center for Fruit Crops at the Lake of Constance (KOB) (Supplementary Materials and Methods S6). It sampled for fourteen days on the same piece of plastic film. Consecutive samples were collected (one sample per fourteen days) from 1 February to 30 June. In 2020, a Mycotrap was placed on a similar leaf litter deposit at the Research Institute of Organic Agriculture FiBL (Frick, Switzerland) and sampled from 1 March to 30 June onto Vaseline coated plastic film. The leaf litter for this deposit was collected in the Rickenbach orchard on 28 February 2020. Spore numbers in the samples were quantified by TaqMan® qPCR as previously described.

We hypothesized that beside leaf litter other inoculum sources, i.e. bark, bud or fruits, might be relevant for disease outbreaks. On 28 February 2020, we collected bark and bud samples from six ‘Otava’ and six ‘Rubinola’ trees in the Rickenbach orchard and on 17 March 2020 from six ‘Topaz’ trees in the KOB orchard. Bark samples were collected as described by Arrigoni et al. (2018). Briefly, bark curls of 20 mm length, 5 mm width and 1 mm thickness were taken using a flame-sterilized scalpel. Each bark sample consisted of a pool of 30 bark curls from a single tree’s upper trunk and lower branches. Ten terminal buds per tree were collected. We further searched for fruit mummies hanging in the trees in the Rickenbach orchard on 28 February 2020. Five pieces of the skin of each mummy (with a diameter of about 2 to 4 mm and a thickness of 2 to 5 mm) were pooled. Samples were lyophilized and ground in a mixer mill MM 200 (Retsch GmbH, Haan, Germany) at 30 Hz for 30 s. A sample of 100 mg of the ground powder (or as much as was available) was subject to DNA extraction using a NucleoSpin™ DNA Stool Kit (Macherey-Nagel, Oensingen, Switzerland). We found only one sample of overwintered leaves hanging in a tree. This sample was processed the same way but without prior lyophilization. Samples were assessed for the presence of *Dc* DNA by the TaqMan® qPCR assay described above.

Finally, to assess whether the identified DNA on bark, buds and fruit mummies originates from viable and infectious fungal tissues, we tried to infect apple leaves in the laboratory from *Dc* PCR positive bud, bark, fruit samples. 400 to 500 mg of fruit, bark, and bud tissue were cut into small pieces and wetted with 3.5 mL of Volvic® natural spring water (n=3). Each of the samples was distributed onto three apple leaves. The experiment was performed twice, once with container grown ‘Topaz’ trees and once with ‘Topaz’ apple seedlings. As controls, three apple leaves were incubated with laboratory infected apple leaves with *Dc* acervuli (positive control) and healthy apple leaves (negative control) processed the same way as described for bark, bud, and fruit samples. In addition, three leaves were inoculated with pieces of a *Dc* positive leaf that was found in the tree crown after winter. The plants were incubated at 20°C with a 16/ 8 h day/night cycle. The first four days the relative humidity was 100 %, followed by 50 % for the rest of the experiment. Symptoms were assessed after three weeks. Leaves showing any sort of spots or necrosis were tested with the *Dc* qPCR.

### Investigation of fruit infections

Apples with symptoms of AB were collected in September of 2020 and 2021 and stored at 4°C until March the following year, when acervuli had developed. The acervuli were examined under a stereo microscope M205C (Leica Microsystems Switzerland, Heerbrugg, Switzerland). The apple tissue containing acervuli were stained with cotton blue in lactic acid and examined with a Leica DM2000 LED microscope (Leica Microsystems Switzerland, Heerbrugg, Switzerland). Images of the apple tissue containing acervuli were captured using a Jenoptic Gryphax Subra camera (Jenoptic AG, Jena, Germany). Conidia formed on cool-stored fruits were tested for their infectivity to apple leaves. To this end, necrotic lesions with acervuli were cut and added to 1 mL Volvic ® natural spring water in 2 mL Eppendorf tubes, vortexed for 30 s and the resulting spore suspension was used to inoculate leaves of six ‘Topaz’ seedlings. Twelve drops of 5 μL suspension were pipetted onto two leaves per seedling. The plants were incubated as described above for infection experiments with bark, bud and fruit mummies.

## RESULTS

### New qPCR allows quantification of *D. coronariae* conidia

To quantify *Dc* conidia in spore trap samples, we developed a new TaqMan® qPCR method with the primers and probes Dc_09 (Table 1). Different primers and hydrolysis probes targeting the ITS gene of *Dc* were tested. Among them, primers and probe Dc_09 performed best (E= 94.2 %, R^2^= 0.995) also when compared to the published primer Mc_ITS (Oberhänsli et al. 2014) (E= 83 %, R^2^ = 0.88). Specificity tests for primer Dc_09 revealed sound amplification of DNA from different isolates of *Dc* (Fig. 2A), but neither amplification with DNA of other fungal species nor with apple DNA (Fig. 2A, Supplementary Table S4).

**Fig. 2.**
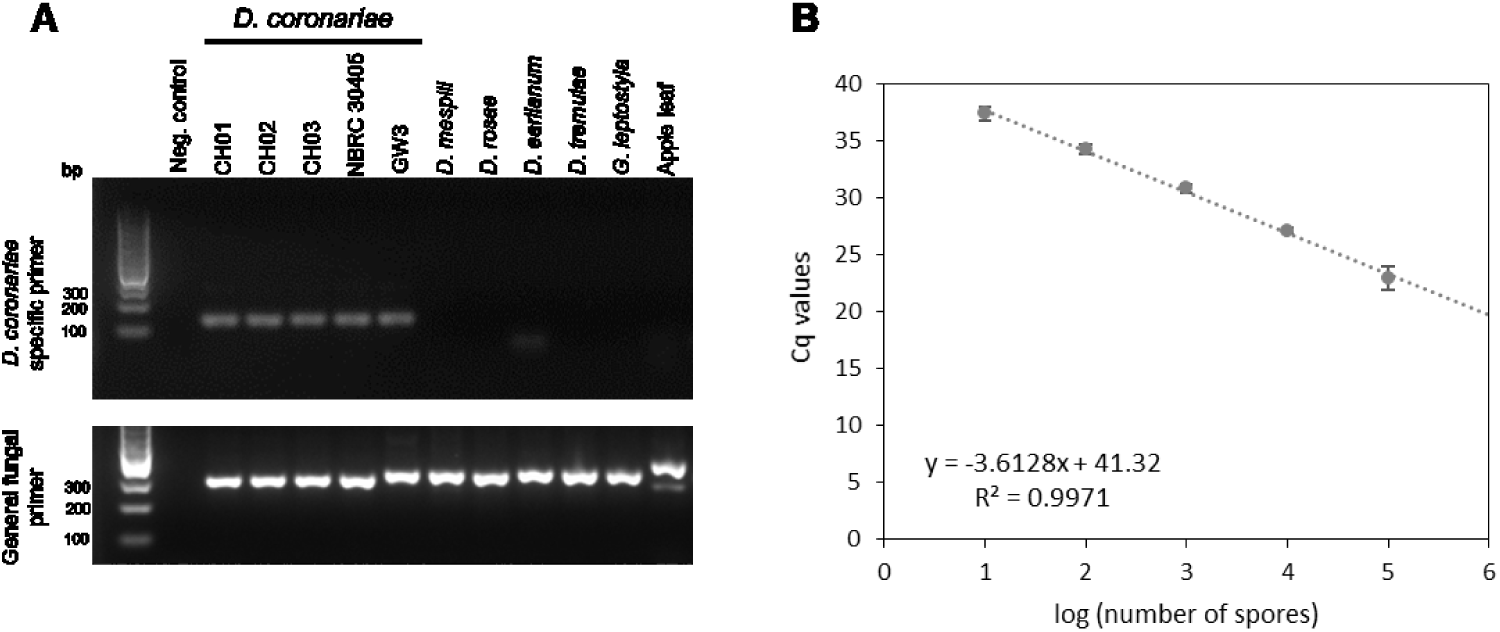
*Diplocarpon coronariae* specific qPCR. **A)** Specificity test of Primer Dc_09. A PCR was performed with DNA from different *D. coronariae* isolates, related *Diplocarpon* species (*D. mespili, D. rosae, D. earlianum*), and with DNA of other pathogens causing Marssonina diseases (*Drepanopeziza tremulae* and *Gnomonia leptostyla*). A 2 % TAE agaraose gel stained with ROTI® GelStainRed (the ladder marker is a peqGOLD O’range 100bp DNA –Ladder (peqlab) **B)** *In vivo* standard curve for calculation of spore numbers. A volume corresponding to 10^5^, 10^4^, 10^3^, 10^2^ and 10^1^ conidia was pipetted onto a Vaseline coated plastic film and DNA was extracted the same way as was done from the Mycotrap and rotating arm spore traps, i.e. using a ZymoBIOMICS® Quick-DNA Fungal/Bacterial Microprep Kit. Samples were spiked with an APA9 plasmid for normalization of DNA extraction and normalized Cq values are depicted. Error bars represent the standard deviation of the mean of seven qPCR runs. The mean efficiency was 89.1 %.

The amplification of the qPCR for conidial counts on Vaseline coated plastic film, which was the spore trapping surface used for field samples, exhibited a linear response with an efficiency of 89.1 % (R^2^ = 0.997) (average of seven qPCR runs) (Fig. 2B). The qPCR assay allowed the consistent detection of as few as ten conidia; however, the amplification was unreliable with three conidia per sample. Therefore, the limit of quantification is considered to be 3 to 10 conidia.

The SYBR Green based hydrolysis probe assay revealed similar sensitivity and specificity as the TaqMan assay (Supplementary Fig. S1). However, we used the TaqMan qPCR for our field experiments since it allowed multiplexing of the *Dc* specific qPCR with the APA9 specific qPCR (plasmid standard), which we used to normalize the *Dc* quantification data with the DNA extraction efficiency.

### Apple blotch disease progress in two Central European apple orchards

At the Rickenbach field site and at the KOB field site disease progression on ‘Topaz’ started with the appearance of first signs of the pathogen (i.e. visible formation of acervuli) and symptoms of AB (yellowing of the leaf), which varied between years, but occurred at the latest in August in both orchards and all years. Progress of AB was generally characterized by a steady increase in severity. However, differences in disease severity were observed between years. At KOB, for example, AB was more severe in 2017 compared to other years (Fig. 1A). Rainfall was more frequent from June through October in 2017, i.e. 78 days of rainfall compared to between 36 and 57 days in the other years (Supplementary Fig. S2). At both sites, 2018 was characterized by an exceptionally hot and dry summer (only 36 and 33 days with rain from June to the end of September at KOB and Rickenbach, respectively) (Supplementary Figs. S2, S3) resulting in lower severity of AB (Fig. 1A). At Rickenbach, AB was most severe in 2019 and 2020; which was not the case at KOB (Fig. 1A).

At the Rickenbach site we identified ‘Otava’, ‘Rajka’, ‘Florina’, and ‘Topaz’ as apple cultivars highly susceptible to AB (Figs. 1B, C). In contrast, ‘Bohnapfel’, ‘Blauacher’, ‘Tobiässler’, and ‘Schneider’ were relatively tolerant with low AB severity, even at the end of the season (Fig. 1C). ‘Rubinola’, had no or very low severity, despite a high inoculum pressure in the orchard (Figs. 1B, C).

### Monitoring dispersal of *Dc* conidia in the field

In 2017 and 2018, the first clearly visible symptoms on ‘Otava’ trees developed by mid-June and early July, respectively. Therefore, considering an incubation period of about three weeks (Lee et al. 2011), the first infections were expected to occur towards late May.

In 2019, very large numbers of spores were detected by the Mycotrap on the ground during the first half of May (Fig. 3A). Subsequently, spore numbers generally decreased, except for a peak in the first week of June. In May, leaves with spotting indicative of AB were collected and tested for *Dc* by qPCR, revealing the first *Dc* positive leaf on 17 May. The first unambiguous symptoms were observed three weeks later, on 7 June (Fig. 3A). The next spore peak was detected on 24 June after a rainy period. At that time, ‘Otava’ trees already showed severe leaf yellowing, indicating the emergence of secondary inoculum. In July, large spore numbers were recorded in the air on most days. The second Mycotrap in the crown of an ‘Otava’ tree detected more spores than the Mycotrap on the ground, except on 24 and 25 June (Supplementary Fig. S4).

**Fig. 3.**
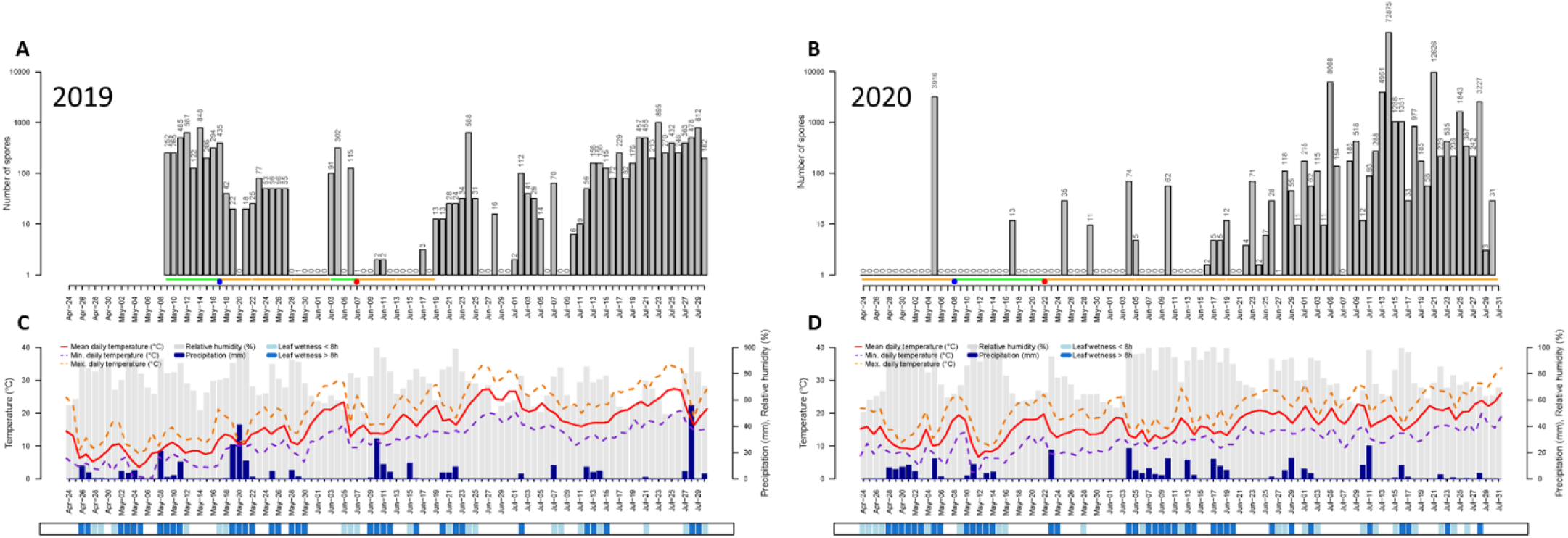
*Diplocarpon coronariae* (*Dc*) spore dispersal in 2019 **(A)** and 2020 **(B)**.Daily spore numbers of *Dc* captured by a Mycotrap placed within an ‘Otava’ tree row of apple trees. The number of spores was calculated based on normalized Cq values from the TaqMan qPCR assay with primer Dc_09 using a standard curve with known numbers of conidia. The green and the yellow lines indicate the periods when a bait plant was exposed in the orchard next to the Mycotrap and whether the bait plant developed AB symptoms (yellow) or not (green). The two blue circles indicate days where first leaves with ambiguous AB symptoms tested positive for *Dc*. The red circles indicate first unambiguous AB symptoms in the orchard. **Weather** data were obtained fromAgrometeo (location “Liebensberg”) **(C, D)** including temperature (°C), relative humidity (%), precipitation (mm), and leaf wetness (light blue, <8h; dark blue, >8h).

The first apparent symptoms on bait plants developed between 17 and 22 May (Fig. 3A). From 22 May on, at least one bait plant showed typical AB symptoms of leaf yellowing, except for the period from 3 to 7 June. The most severe leaf yellowing and leaf drop were observed on the bait plants exposed between 7 and 13 June, followed by those exposed between 13 and 19 June.

In 2020, very few spores were captured by the trap on the ground before bud break in March (Fig. 4). The first peak in spore production was captured on 5 May by the trap on the ground at the end of a rainy period that had followed an extremely dry April (Figs. 3B, 4, Supplementary Fig. S2). The bait plants standing in the orchard from 24 April to 8 May were the first to develop AB symptoms. The first *Dc* positive leaf based on a qPCR test was on 8 May. Leaves with clear symptoms and production of conidia were observed on 22 May 2020, which was approximately two weeks earlier than observed in 2019.

**Fig. 4.**
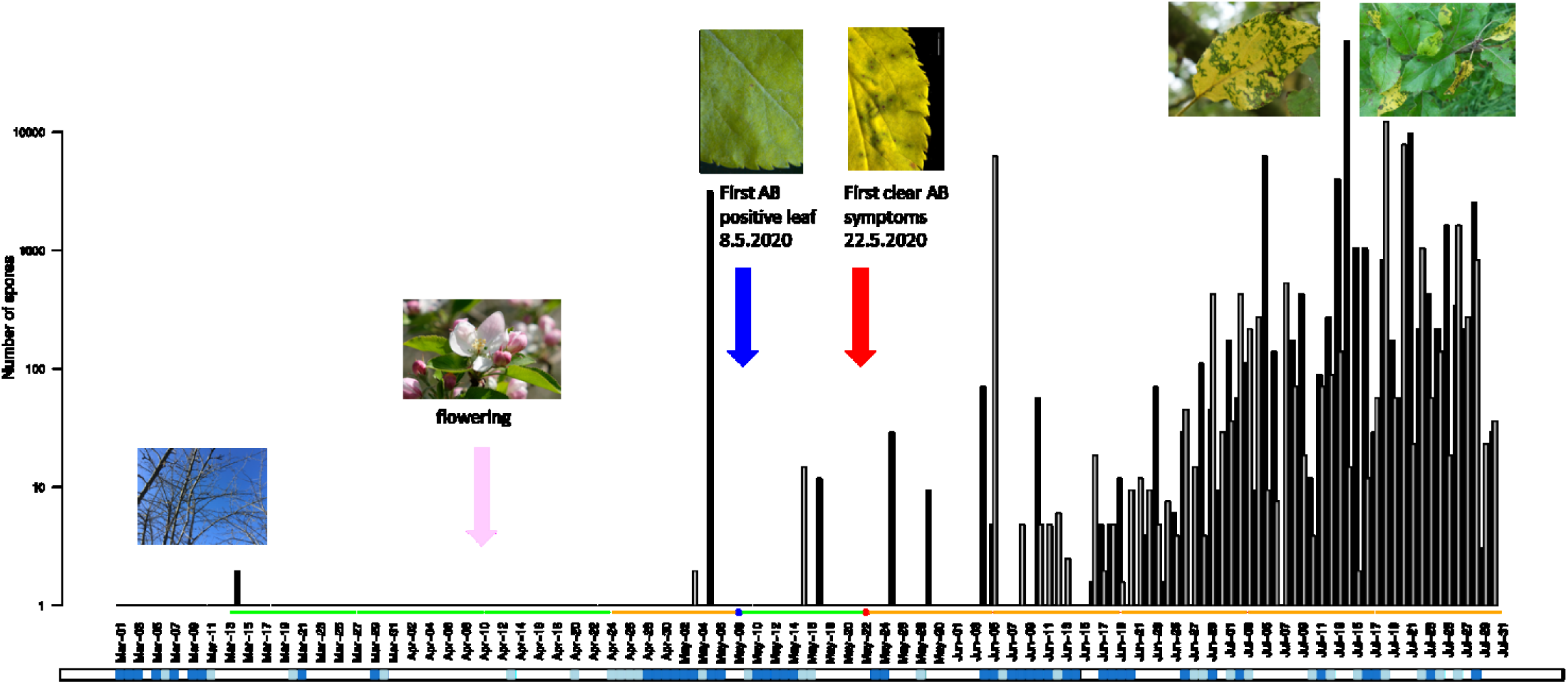
*Diplocarpon coronariae* (*Dc*) spore dispersal on the ground and in the tree crown in 2020. Daily spore numbers of *Dc* captured by the Mycotraps placed on the ground within an ‘Otava’ tree row of apple trees (black bars) and within the tree canopy (grey bars). The number of spores was calculated based on normalized Cq values from a TaqMan qPCR assay with primer Dc_09 using a standard curve with known amounts of conidia. The green and the yellow lines indicate the periods where a bait plant was exposed in the orchard next to the Mycotrap and whether the bait trap plants developed apple blotch symptoms (yellow) or not (green). Estimated duration of leaf wetness is indicated as light blue (<8h) and dark blue (>8h) bands.

A second peak in spore production was detected on 5 June by the spore trap in the tree crown. As in 2019, spores were detected in the air almost every day by both traps from mid-June to the end of July (Figs. 3, 4). Numbers of spores varied, but no significant correlation was found between spore numbers and precipitation (Supplementary Fig. 6). In Mycotrap samples in early June (lower trap: 5 to 19 June, upper trap: 31 May to 9 June), we experienced technical problems resulting in poor DNA extraction efficiency. In cases with signal by the *Dc* qPCR, the low efficiency could be taken into account by normalization with the internal plasmid standard. However, in cases where there was no signal *Dc*, we were unable to conclude whether the zero value was due to no spores or the low efficiency precluding detection. Thus, for some days during the first half of June, *Dc* spores might have been present in the air, although no catch is indicated (Figs. 3B, 4).

In 2020, all bait plants placed in the orchard after 22 May developed AB symptoms. From 22 May to 5 June, bait plants exhibited approximately 50 to 80% of leaves symptomatic, and after 5 June incidence of leaves exhibiting AB exceeded 90% (Supplementary Fig. S5).

### Dispersal of conidia of *Dc* with distance from an inoculum source

Rotating-arm spore traps detected the first spores after 22 May in 2019 and 2020. Spore numbers detected in rotating-arm spore traps were generally lower than those captured in the Mycotrap samples (Table 2, Figs. 3, 4). The highest spore numbers were generally detected in traps on ‘Otava’ trees, which exhibited severe symptoms of AB, while almost no spores were detected in traps on ‘Rubinola’ trees. Despite being only two rows (18 m) apart from highly diseased trees, the ‘Rubinola’ trees were virtually free of AB, although some leaf yellowing and premature leaf fall was observed on ‘Rubinola’ trees. The symptoms on ‘Rubinola’ were distinctly different from typical AB, and when such leaves were tested by qPCR no *Dc* DNA was detectable (n = 7). Furthermore, no spores were detected in the trap 50 m beyond the orchard edge with one exception: during the period from 22 May to 3 June 2019 spores of *Dc* were detected under ‘Rubinola’ trees and outside the orchard. A strong wind event with wind speed up to 37 km/h combined with rain on 21 May may have transported some infected leaves towards the ‘Rubinola’ trees and outside the orchard explaining these results.

**TABLE 2.**
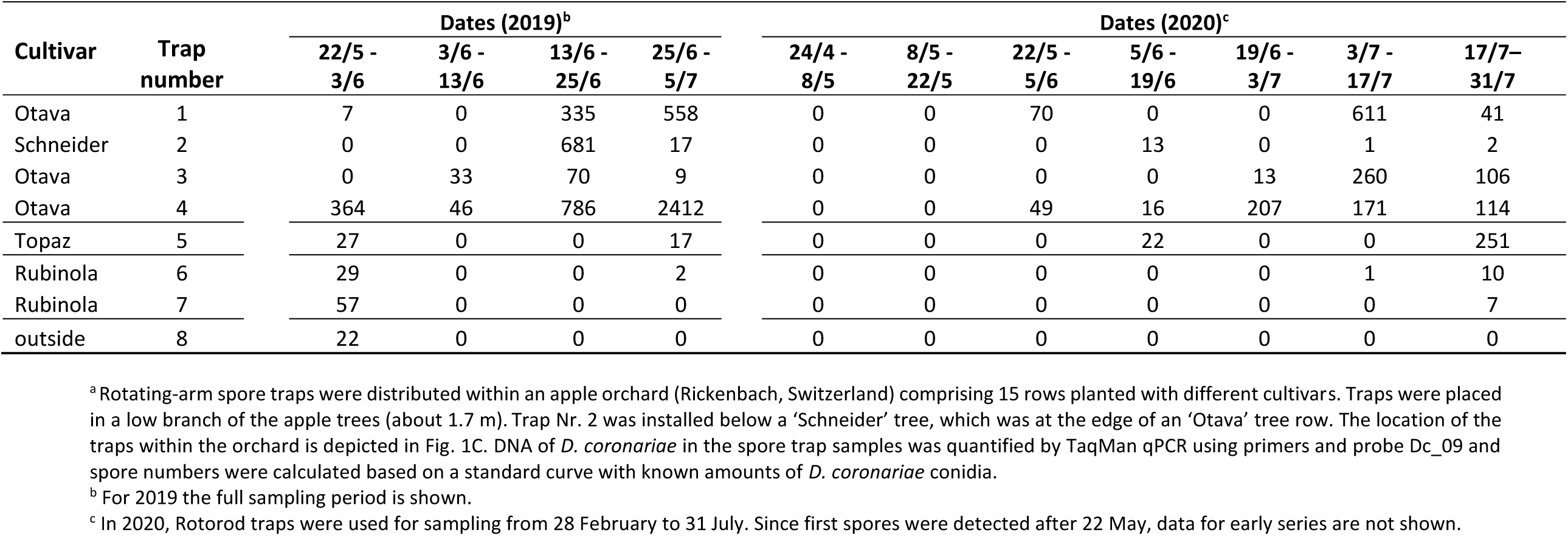
Numbers of conidia of *Diplocarpon coronariae* captured using rotating-arm spore traps in an apple orchard in Switzerland.^a^

Besides the dispersal of *Dc* spores via the air, rainwater splashes might be an important factor for spreading the disease within a tree. Therefore, we collected rainwater below highly infected trees (‘Otava’, ‘Topaz’), below symptomless trees (‘Rubinola’), between the rows under the open sky and outside the orchard during a rainy period in September 2020. We detected several hundred spores per mL rainwater below ‘Otava’ trees, fewer spores between rows, and almost no spores below non-infected ‘Rubinola’ trees or outside the orchard (Supplementary Fig. S6).

### Origin of primary infections

At KOB in 2019, the first spores were released from a deposit of overwintered leaf litter in the second half of April (Table 3). At the Research Institute of Organic Agriculture FiBL in 2020, however, a few spores were detected above a leaf litter deposit in the second half of March. In 2020, no spores were detected in April at FiBL, which was characterized by extraordinarily dry weather (Supplementary Fig. S3), but the spores were again released from the leaf litter deposit during May (Table 3). Spores were detected in the second half of June at KOB and FiBL. Generally, spore numbers detected in the experiments were very low and thus not reliably quantifiable (LOQ = 10 spores).

**TABLE 3.**
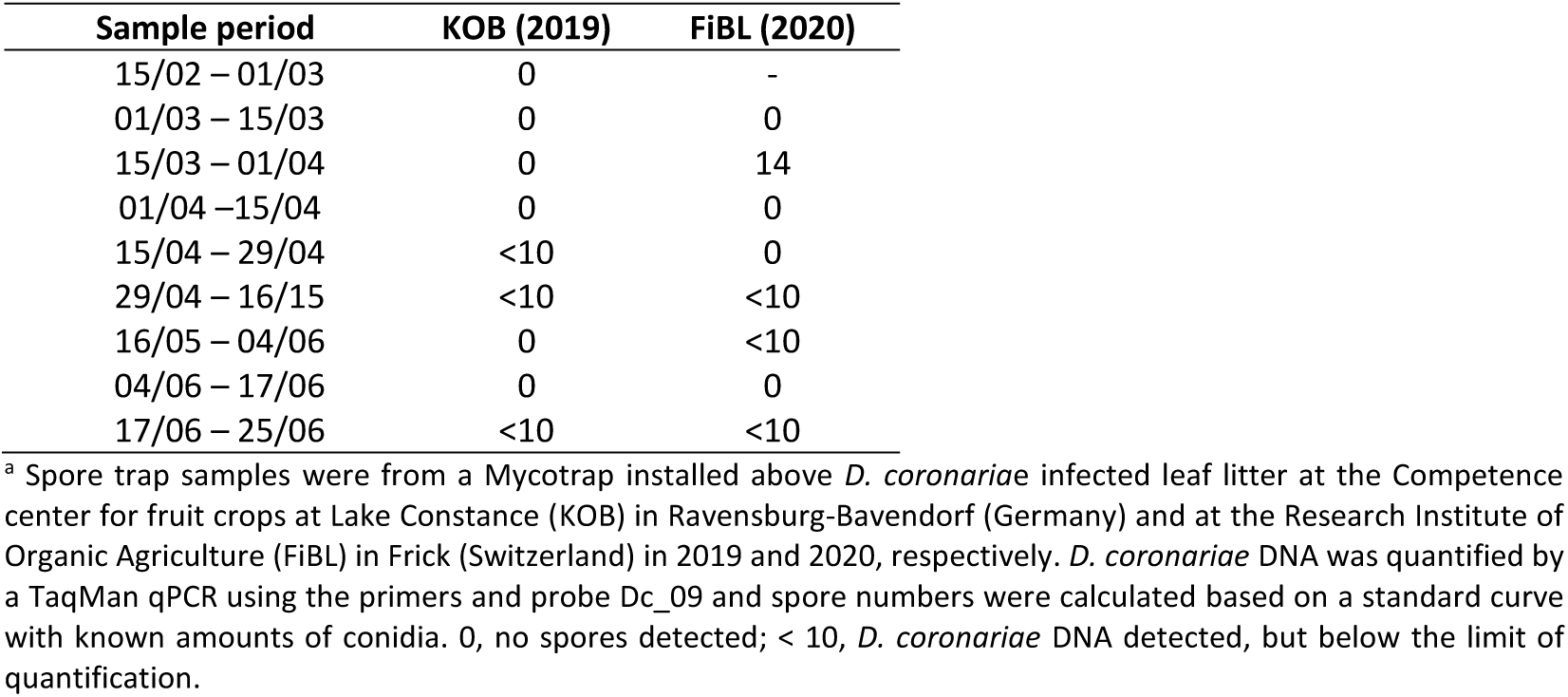
The number of conidia of *Diplocarpon coronariae* collected in a Mycotrap spore trap above leaf litter on an orchard floor in 2019 and 2020.^a^

| Sample period | KOB (2019) | FiBL (2020) |
| --- | --- | --- |
| 15/02 – 01/03 | 0 | - |
| 01/03 – 15/03 | 0 | 0 |
| 15/03 – 01/04 | 0 | 14 |
| 01/04 – 15/04 | 0 | 0 |
| 15/04 – 29/04 | <10 | 0 |
| 29/04 – 16/05 | <10 | <10 |
| 16/05 – 04/06 | 0 | <10 |
| 04/06 – 17/06 | 0 | 0 |
| 17/06 – 25/06 | <10 | <10 |

<sup>a</sup> Spore trap samples were from a Mycotrap installed above *D. coronariae* infected leaf litter at the Competence center for fruit crops at Lake Constance (KOB) in Ravensburg-Bavendorf (Germany) and at the Research Institute of Organic Agriculture (FiBL) in Frick (Switzerland) in 2019 and 2020, respectively. *D. coronariae* DNA was quantified by a TaqMan qPCR using the primers and probe Dc\_09 and spore numbers were calculated based on a standard curve with known amounts of conidia. 0, no spores detected; < 10, *D. coronariae* DNA detected, but below the limit of quantification.

While all bark and bud samples of ‘Rubinola’ were negative for *Dc*, the majority of bark and bud samples from ‘Topaz’ and ‘Otava’ trees were tested positive for *Dc* (Table 4). Bark samples exhibited high Cq values close to the limit of detection corresponding to low amounts of DNA. Bud samples contained up to 10^2^ to 10^4^ times higher levels of *Dc* DNA than bark samples. The highest amount of *Dc* DNA was detected in a bud sample of an Otava tree with a Cq value of 25.6 (Table 4), which corresponds to the DNA of 10^4^ conidia.

**TABLE 4.** Detection of *Diplocarpon coronariae* on the bark and in the buds of apple trees during winter. The apple trees had symptoms of apple blotch the previous summer.

| Apple organ | Field site<sup>a</sup> | Apple cultivar<sup>b</sup> | Tree number<sup>c</sup> | Normalized C<sub>q</sub> value<sup>d</sup> |
| --- | --- | --- | --- | --- |
| Bark | KOB | Topaz | 18-17 | 38.7 |
| Bark | KOB | Topaz | 18-25 | 41.1 |
| Bark | KOB | Topaz | 18-37 | ND |
| Bark | KOB | Topaz | 18-38 | 42.3 |
| Bark | KOB | Topaz | 18-39 | 42.0 |
| Bark | KOB | Topaz | 18-40 | ND |
| Bark | Rickenbach | Otava | 4-10 | 41.1 |
| Bark | Rickenbach | Otava | 4-11 | ND |
| Bark | Rickenbach | Otava | 4-13 | 39.9 |
| Bark | Rickenbach | Otava | 4-14 | 40.71 |
| Bark | Rickenbach | Otava | 4-15 | 38.10 |
| Bark | Rickenbach | Otava | 4-16 | 38.30 |
| Bark | Rickenbach | Rubinola | 10-4 | ND |
| Bark | Rickenbach | Rubinola | 10-5 | ND |
| Bark | Rickenbach | Rubinola | 10-6 | ND |
| Bark | Rickenbach | Rubinola | 10-7 | ND |
| Bark | Rickenbach | Rubinola | 10-8 | ND |
| Bark | Rickenbach | Rubinola | 10-9 | ND |
| Bud | KOB | Topaz | 18-17 | ND |
| Bud | KOB | Topaz | 18-25 | 36.3 |
| Bud | KOB | Topaz | 18-37 | 28.0 |
| Bud | KOB | Topaz | 18-38 | 31.4 |
| Bud | KOB | Topaz | 18-39 | 36.1 |
| Bud | KOB | Topaz | 18-40 | 37.5 |
| Bud | Rickenbach | Otava | 4-10 | 33.8 |
| Bud | Rickenbach | Otava | 4-11 | ND |
| Bud | Rickenbach | Otava | 4-13 | 41.6 |
| Bud | Rickenbach | Otava | 4-14 | 33.8 |
| Bud | Rickenbach | Otava | 4-15 | 25.6 |
| Bud | Rickenbach | Otava | 4-16 | 30.7 |
| Bud | Rickenbach | Rubinola | 10-4 | ND |
| Bud | Rickenbach | Rubinola | 10-5 | ND |
| Bud | Rickenbach | Rubinola | 10-6 | ND |
| Bud | Rickenbach | Rubinola | 10-7 | ND |
| Bud | Rickenbach | Rubinola | 10-8 | ND |
| Bud | Rickenbach | Rubinola | 10-9 | ND |

<sup>a</sup> Bud and bark samples were collected from apple trees in the Rickenbach orchard (Switzerland) on 28 February and on 13 March 2020 and from trees in the KOB orchard (Germany) on 17 March 2020.
<sup>b</sup> 'Topaz' and 'Otava' trees were heavily diseased with apple blotch in 2019, while 'Rubinola' trees showed no symptoms.
<sup>c</sup> The tree number indicates the position of the tree within the orchard and consists of row number and tree number within row.
<sup>d</sup> C<sub>q</sub> (quantification cycle) values result from a TaqMan qPCR with primers and probe Dc\_09. The APA9 plasmid was used to normalize for DNA extraction efficiency. ND, not detected.

In spring 2020, four out of five fruit mummies, which were collected from trees with strong AB infestation in 2019, were tested slightly positive (Cq values between 33.65 and 40.38) (Fig. 5C, Supplementary Table S5), while fruit mummies from trees without AB symptoms the previous year were negative (Supplementary Table S5). For comparison, a leaf sample still hanging in a susceptible ‘Florina’ tree after the winter of 2019 was *Dc* positive (Cq, 25.83).

**Fig. 5.**
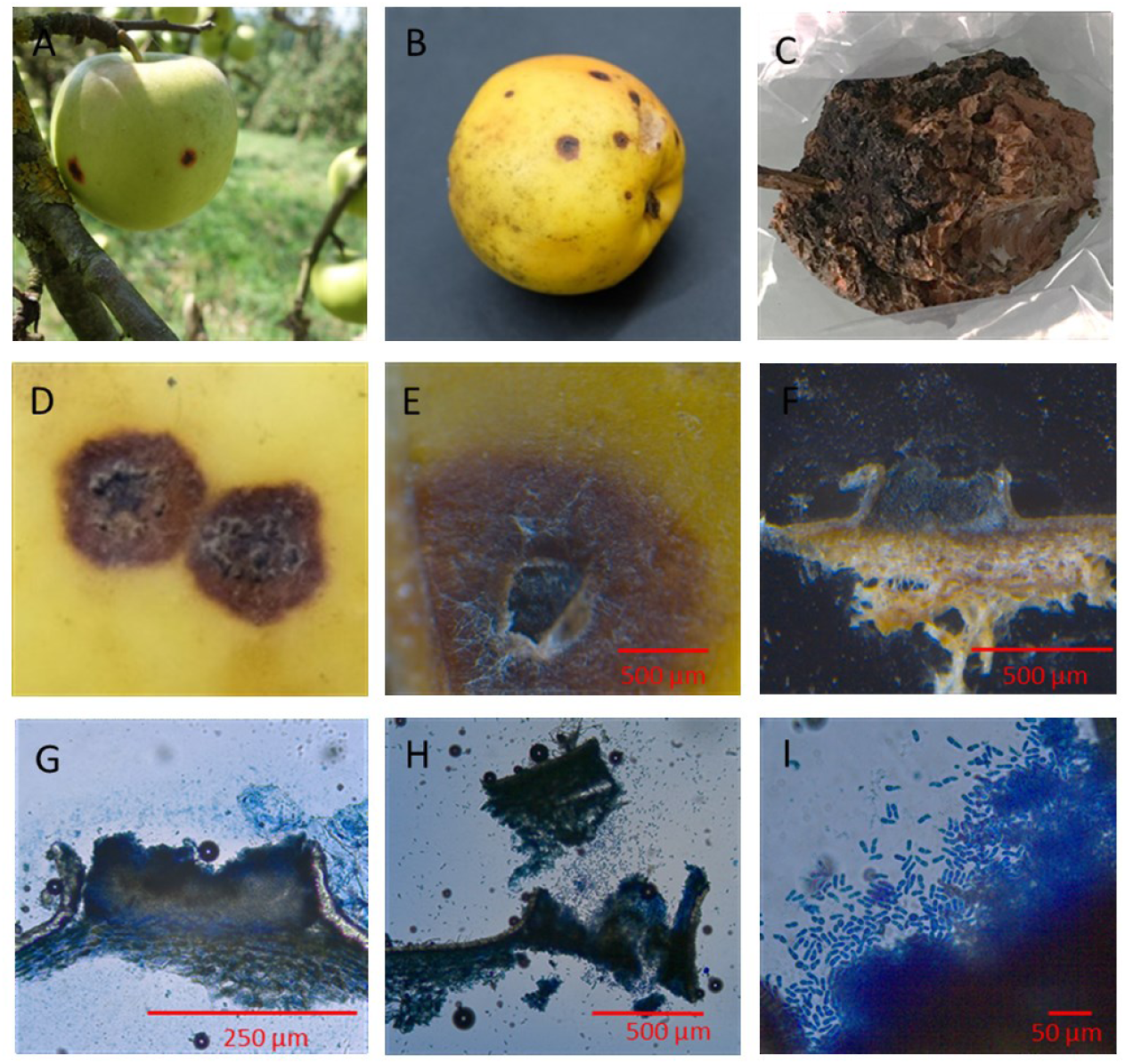
Fruit infections with *Diplocarpon coronariae (Dc)*. A) ‘Otava’ apple with apple blotch symptoms hanging in a tree in September 2020 (Rickenbach, Zurich, Switzerland). B) *Dc* infected ‘Otava’ apple after 6 months of storage at 4°C. C) ‘Florina’ fruit mummy found hanging in a tree in February 2020, and tested positive for *Dc* by TaqMan qPCR. D-I) Acervuli and conidia developed on *Dc* infected ‘Otava’ apple collected in the field in September 2019 and stored at 4°C for six months. F-G) Longitudinal section through an acervulus. H) Conidia released from an acervulus. I) *Dc* conidia. D-F) Pictures were taken with a stereo microscope M205C (Leica Microsystems Switzerland, Heerbrugg, Switzerland). G-I) Samples were stained with cotton blue in lactic acid and pictures were taken with a Leica DM2000 LED microscope (Leica Microsystems Switzerland, Heerbrugg, Switzerland).

Attempts to infect apple leaves in the laboratory by applying wetted pieces of bark, buds, fruit mummies and leaves from previously AB symptomatic trees did not result in successful infections (data not shown). But *Dc* infected fruits can serve as an inoculum source for infection of leaves after six months of storage. In fall 2020 and 2021, *Dc* infections were observed on ‘Otava’ fruits in the Rickenbach orchard (Fig. 5A). In the following spring, after storing fruits at 4°C for six months, conidia formed (Figs. 5B, D-I), and the conidia infected the leaves of apple seedlings in the laboratory (Supplementary Fig. S7).

## DISCUSSION

### Development of a *D. coronariae* specific qPCR

To date, spore trap catches of *Dc* conidia have been quantified using microscopy (Kim et al. 2019). However, microscopy is labor intensive and identification of the *Dc* spores in field samples may be challenging. We developed a qPCR method to provide a very sensitive means for quantification of as few as ten conidia per sample, which is in a similar range to qPCRs developed for other fungi (Calderon et al. 2002; Dvořák et al. 2015; Luchi et al. 2013). The qPCR method further allows diagnosis of AB prior to developing unambiguous visual symptoms. Early diagnosis is especially useful when small necrotic lesions appear that are indistinguishable from symptoms from other causes. In summary, we established a new qPCR method, which allowed the quantification of *Dc* spores in spore trap samples, and that can be used to detect initial infections in the field, and that enabled the identification of *Dc* DNA in various apple organs.

qPCR, though it requires several processing steps, is a rather fast method and allows the processing of many samples concurrently. However, it has the disadvantage to only allow conclusions regarding the presence of DNA, but not regarding the fungal organ (e.g. mycelium, ascospore, conidia, spermatia) or its viability and infectious potential. Therefore, visual assessments and infection experiments are needed to understand how *Dc* exists on bark, bud and fruit samples and the role of the tree organs in the fungal life cycle.

### Timing of primary infections

To determine the timing of primary infections by *Dc*, we combined data on airborne spore catches with symptom development both in the field and on a consecutive series of exposed bait plants. In the first three study years (2016-2018), clear AB symptoms were observed, the earliest showing in June. However, testing leaves with ambiguous necrotic spots by qPCR in 2019 and 2020 revealed that the first leaves of highly susceptible varieties were already infected with *Dc* in May (17 May 2019 and 8 May 2020). The infections were most likely caused by spores released in late April or early May. The first spore peaks were detected in the orchard in early May in both 2019 and 2020. In 2019, spore traps were installed on 9 May 2019. Thus, it is possible that spores were already present one or two weeks earlier. In 2020, the experiment was initiated at the end of February, and the first clear spore peak was detected by the Mycotrap on the ground on 5 May. We conclude that the spores collected in early May likely caused the first infections in the field since the first bait plants in the field that developed symptoms were exposed from 24 April to 8 May 2020.

In line with the results of the spore traps in the orchard, the spore traps located above leaf litter caught conidia of *Dc* at the end of April and early May in both study years. The few spores detected above the leaf litter and in the field in March 2020 were released before bud break, and we, therefore, conclude that they were irrelevant to primary leaf infections.

Our qPCR method does not distinguish between conidia and ascospores, but apothecia of *Dc* have not yet been observed in Europe (Wöhner and Emeriewen 2019), indicating it is unlikely they were the spore type captured. Although not yet observed, the importance of ascospores as primary inoculum in Europe should be subject to further research. A recent population genetic study suggested a mixed sexual and asexual reproduction of *Dc* in Europe, although *Dc* populations in Europe are genetically homogenous, clonal and dominated by a few multi-locus genotypes (Oberhänsli et al. 2021).

The first spore peaks of the season were observed at the end of, or shortly after rainy periods, whereas no spores were detected during dry periods before rain events, e.g. April 2020. The result indicates that rain might be required for the release of conidia of *Dc* from leaf litter, and acervuli might need a wetting period prior to release of conidia. This hypothesis may explain the slightly negative correlation found for May 2019 between daily spore catches and precipitation (Supplementary Table S6).

Based on our results, primary infections of apple with *Dc* occur in Central Europe in late April or early May after periods of rain, which is earlier than was previously assumed (Sutton et al. 2014). Moreover, the spore catches in March indicate that *Dc* spore discharge may occur before April and, depending on weather conditions and phenological stage of the host, infections could be possible. Knowing the timing of primary infections can be valuable as a basis for management actions to preventing epidemics of AB. While protection strategies against AB in Europe are currently focused on the summer months (Hinrichs-Berger 2015), prevention of primary infections in the early spring might be an important additional step for more efficacious management strategies in the future.

### Leaf litter and other sources of primary inoculum

In addition to the timing of primary infections, we were also interested in the origin of the primary inoculum. Several studies report that conidia inside acervuli overwinter in leaf litter (Back et al. 2015; Dong et al. 2015; Gao et al. 2011; Goyal et al. 2018; Lee et al. 2011; Sastrahidayat and Nirwanto 2016; Sharma et al. 2009). This is in accordance with our finding of *Dc* spores above leave litter deposits in spring and also with the observation that in May, most spores were captured in the Mycotrap on the ground, and not in the Mycotrap in the tree canopy.

Small quantities of *Dc* DNA were detectable on fruit mummies hanging in the trees at the end of February, and acervuli with infectious conidia were produced on *Dc* infected fruits after overwintering at 4°C. Moreover, considerable quantities of *Dc* DNA were detected on buds, and small amounts of *Dc* DNA were found on the bark of trees that had AB the previous season. All these organs are reported as overwintering sites for other apple pathogens, including *Monilia* spp. on fruit mummies, *Podosphaera leucotricha* in buds and *Neofabraea* spp. on bark (Sutton et al. 2014). Moreover, *D. rosae*, a close relative of *Dc* causing black spot disease on roses, overwinters, in addition to leaf litter, on bud scales and stems (Cook 1981). To date, it is unclear what the survival structures of *Dc* on bark and buds are (e.g. mycelium, spores), and whether they are infectious. The fact that we could not cause leaf infections from bud, bark or fruit mummy tissues may be due to technical issues, and the role of the various apple tree organs for overwintering of *Dc* requires further investigation. *Dc* present on buds and wood in propagation material could be exchanged between nurseries across Europe and may explain the rapid spread of this disease to many European apple production areas since its first detection in Northern Italy (Oberhänsli et al. 2021). Our observation that conidia produced on fruits after six months in storage at 4°C are able to infect apple leaves provides an indication that fruit-derived conidia might represent an additional source of primary inoculum at least on a local scale. Moreover, our description of fruit infection adds to our very limited knowledge regarding the role of fruit infections in the epidemiology of AB (Wöhner and Emeriewen 2019).

### Timing of secondary infections and epidemic development

By the end of May (2020) or the beginning of June (2019), the first leaves that were producing acervuli and conidia of *Dc* were observed, initiating the secondary phase of infection. In contrast to the first major peak of conidia captured in May 2020, the second major peak (5 June) was not detected by the Mycotrap on the ground but by the Mycotrap situated in the tree canopy. We hypothesize that the conidia produced in early May originated from overwintered leaf litter, resulting in higher spore counts in the Mycotrap on the ground. But in June, the conidia originated from the recently infected leaves in the tree canopy, resulting in a higher spore count in the Mycotrap situated in the tree canopy. In support of the hypothesis, the rotating-arm spore traps in the tree crown did not capture conidia before the end of May.

From June onwards, damage caused by AB steadily increased in the orchard (Fig. 1A, B). From the middle of June to the end of the experiment, high numbers of spores were detected irrespective of rain events, and no correlation was found between daily precipitation and caught spores in June and July for both years (Supplementary Table S6). This contrasts with our observation of the early spore peaks, which occurred at the end of the rain periods and the observations by Kim et al. (2019). They investigated dispersal of conidia of *Dc* from June to October 2013 and 2015 in Korea and reported spore dispersal peaks during and two days after rain events compared to dry periods. In that study, disease incidence and total spore counts were much lower than we observed at our field site, which might explain some of the differences. Based on our results, we hypothesize that primary inoculum may require water to ripen and discharge, while secondary inoculum are also released on dry days. Lab experiments indicate that *Dc* conidia can survive for several days under dry conditions after landing on a susceptible leaf, and cause infection once the leaf becomes wet (Clémence Boutry et al., unpublished data). If so, all secondary inoculum result in risk of infection whenever leaf wetness occurs.

The development of AB is affected by environmental factors (Li et al. 2011). Rainfall and the subsequent relative humidity are positively correlated with the disease severity in the summer months (Rather et al. 2017b; Sastrahidayat and Nirwanto 2016; Sharma and Sharma 2005). Under controlled conditions, the minimum leaf wetness period for infection is 8 h at 15°C and decreases to 4 h with temperatures between 20 and 25°C (Sharma et al. 2009). Moreover, the incubation period can vary depending on the weather conditions. At a temperature range of 15 to 25°C and seven days of leaf wetness, the incubation period is 10 to 20 days (Harada et al. 1974). In our study, the bait plants showed symptoms within one to four weeks after return from the orchard, with a shorter incubation period for bait plants placed in the orchard later in the season (June, July) compared to the beginning of the season (May). In 2020, the first necrotic spots indicative of AB and the more developed symptoms were observed earlier and more rapidly than in 2019. The faster disease progression in late spring 2020 was likely due to higher May temperatures compared to the temperatures in 2019.

### Spore dispersal in relation to distance from an inoculum source

Rotating-arm spore traps placed in the orchard revealed high spore concentrations in AB-diseased trees, but no or few spores in trees without symptoms. The observation could indicate that conidia of *Dc* are not dispersed over large distances. However, as noted there were generally low spore counts captured by the rotating-arm traps compared to the Mycotrap, indicating that the rotating-arm traps were much less efficient in capturing spores in the field (so spores might have been there but the lower sensitivity of the trap precluded detection). Thus, wind might transport some spores over larger distances than we observed. Besides wind dispersal, dispersal in rain splash may contribute to the spread of *Dc* within the tree canopy, as indicated by the very high spore counts in rainwater below highly symptomatic trees (Supplementary Fig. S6).

### *Dc* life cycle

Based on our results we contend that in Europe, the fungus overwinters primarily in infected leaf litter (Fig. 6). However, bark, buds and fruit mummies might be alternative sources of primary inoculum since we detected *Dc* DNA on these organs after winter in previously AB-symptomatic trees. A few spores appear to be released in March, but primary inoculum infections start in late April or early May, depending on the weather conditions. Secondary inoculum spores developed in late May or early June in our field sites, which was within three to four weeks of the primary infections. In May, spore peaks were mainly found at the end of, or after periods of rain; whereas spore load in the air was generally high from the middle of June to the end of July. Thus, every wet period in summer represents a risk for infection. The exact mode of pathogen dispersal has not yet been conclusively determined. Conidia are probably dispersed by wind and rain splash over larger distances; for the spread of *Dc* within a tree canopy or between neighboring tree canopies, rain splash could be a major dispersal agent.

**Fig. 6.**
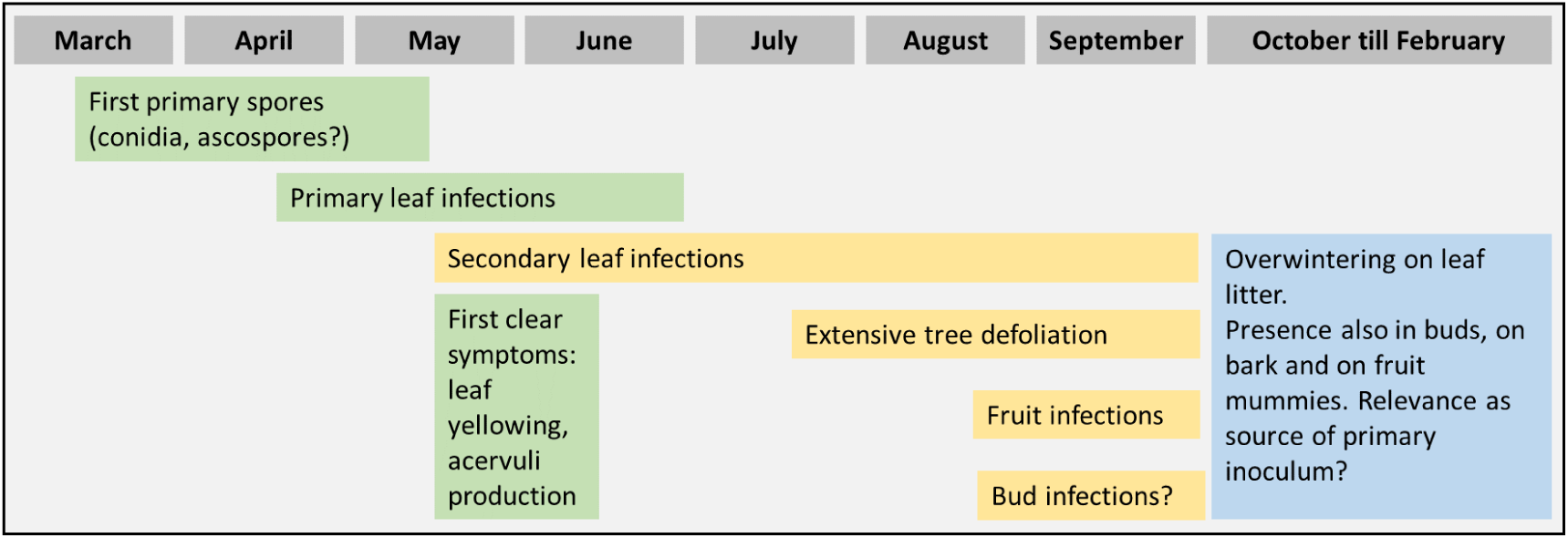
The proposed infection cycle of *Diplocarpon coronariae* (*Dc*) in Central Europe. Based on previous knowledge and the data we presented in the current study, we hypothesize primary spores (conidia since no ascospores have been reported in Europe to date) are released in early spring and cause the first infections between the end of April and the middle of May depending on the weather conditions. Without crop protection, secondary spore inoculum is produced from the end of May onwards. The epidemic of AB spreads and leads to tree defoliation and infection of fruit in severely infested orchards. The fungus overwinters on leaf litter, but is also found on fruit mummies, in buds and on bark after winter. The fruit mummies, buds and bark might represent alternative organs for overwintering. Green, primary infections; yellow, secondary infections; blue, overwintering.

In summary, we developed a sensitive and specific qPCR method to detect and quantify conidia of *Dc* captured in spore traps, which can be combined with the quantification of other apple pathogens, e.g. *Venturia inaequalis* (Meitz-Hopkins et al. 2014), in spore trap samples from apple orchards in the future. Our data contributes to our understanding of AB, especially the timing of early spore dispersal and infections. The information can help to improve disease forecast models for AB, such as the one by RIMpro (RIMpro, Amsterdam, The Netherlands), and provide a basis for early disease management. However, more research is needed to test whether preventing primary infections will reduce disease progress over the season, thereby reducing the need for fungicide applications in the summer. Finally, the presence of *Dc* DNA on bark, buds and fruit mummies indicates that the role of overwintering inoculum on these organs and the anthropogenic dispersal of the pathogen on propagation material should be further investigated.

## ACKNOWLEDGEMENTS

We are very grateful to Jürg and Pascale Strauss, who allowed us to perform major parts of the study in their apple orchard and actively supported us with technical assistance in the field. We thank David Metzger, Mona Blattner, Sumin Bae, Camilla Kappeler, Patrick Widmann, and Adrian Rutzer for their excellent help with fieldwork, disease scoring and laboratory assistance. We are very grateful to Lauren Dietemann for taking the time to review the manuscript for the English. We further thank Monika Maurhofer for valuable scientific input as an examiner on a Master thesis included in this publication. Finally, we are grateful to Marc Trapman for developing the *Dc* infection forecast model ‘RIMPro Marssonina’, which gave important stimuli for our work and has shown to be helpful in disease forecast.

## Notes

### Competing Interest Statement

The authors have declared no competing interest.

