## Supplementary Materials, Tables, and Figures for "Monitoring Spore Dispersal and Early Infections of *Diplocarpon coronariae* Causing Apple Blotch Using Spore Traps and a New qPCR Method"

### Supplementary Materials and Methods

**Supplementary Materials and Methods S1.** Information on spore trap types**.**

#### The seven-day volumetric spore trap (Mycotrap)

The Research Institute Wädenswil (Forschungsanstalt Wädenswil, FAW; current name "Agroscope Wädenswil") developed a volumetric spore trap called Mycotrap (Siegfried et al. 1996), which was rebuilt at the Research Institute of Organic Agriculture (FiBL).

The Mycotrap is an impaction spore trap where the air is sucked in horizontally through a sampling orifice (3 x 15 mm) and impacts a trapping surface on a cylinder inside a chamber. The air suction is achieved by a ventilator as used in airplanes (Micronel D481T-012KA-3, 634 l/min), and the trap has a throughput of 22 L/min. The intake slit is protected from rain by a shield, and a wind vane attached to the head of the device ensures that the opening is always facing the wind so that the collection volume is independent of changes in wind direction. Thanks to a clockwork mechanism, the cylinder slowly rotates (1.66 mm/h resp. 40 mm/d). The cylinder makes a complete turn within seven days, thus allowing for sampling for seven days in a row, but also involves replacing the spore trapping surface after seven days. In our study, the spore trapping surface consisted of transparent plastic film covered with a thin layer of petroleum jelly (Vaseline). The tape was cut into equal daily segments (40 mm, midnight to midnight) and spore DNA extracted from each sampling day separately. Seven-day volumetric spore traps are well suited to monitor the temporal resolution of spore dispersal and allow comparison of spore production with environmental data from the same period (Meitz-Hopkins et al. 2014).

#### The rotating-arm spore trap

Rotating-arm spore traps are available for purchase (Rotorod sampler, Sampling Technologies, Minnetonka, MN) but are often self-designed by researchers at a low cost. A rotating-arm spore trap consists of a U-shaped rod attached to a battery-powered electric motor. The two leading edges of the rod (e.g. 1.1 diameter and 40 mm length, as used in Thiessen et al. (2016)) collect airborne particles by impaction (West et al. 2008). The two leading edges, often made of stainless steel (Choudhury et al. 2017; Klosterman et al. 2014; Kunjeti et al. 2016; Thiessen et al. 2016) or glass (Falacy et al. 2007), are coated with an adhesive, usually silicon grease, applied directly onto the rods or with a tape wrapped around the rods (Calderon et al. 2002). Some innovative solutions for rods are possible, for example, matches covered with double-sided tape (Chandelier et al. 2014). The rods can rotate at different speeds depending on the power of the motor and the battery, e.g. as slow as 30 rpm (Quesada et al. 2018) or as fast as 2400 rpm (Chandelier et al. 2014; Dvořák et al. 2017). The air sampling rate depends mainly on the trapping surface and rotation speed. Therefore, the air sampling rate of rotating-arm spore traps varies greatly and ranges from 20 to 200 liters of air per minute. If the arms rotate fast enough, they suck in air from above and below, similar to fans, so that it is unlikely that they will sample the same air twice, and collection rates close to 100% are theoretically possible (McCartney et al. 1997). The motor and the arms are usually protected from rain by a small shield on top of the spore trap.

The rotating-arm spore trap can be designed light and compact, and due to its low cost, several devices can be deployed in a specific area. This can increase the probability of detection of spores or allow to monitor the distribution of the spores at different heights, directions and distances around a source (Chandelier et al. 2014; Lacey and West 2006). The rotating-arm spore trap is most effective for trapping relatively large fungal spores (>7 µm) because they are most likely to impact, whereas smaller (lighter) particles are likely to be deflected from the spore trapping surface on the leading edges of the arms (Frenz 1999). Therefore, this type of spore trap is not adapted for catching small spores. Dvořák et al. (2015) found that the trapping efficiency for *Hymenoscyphus fraxineus* conidia (size up to 7 x 2.5 µm) tended toward zero. Moreover, rotating-arm spore traps are also more exposed to weather conditions such as heavy rain than other spore traps, and thus spores might get washed off from the trapping surface.

**Supplementary Materials and Methods S2.** Construction of the rotating-arm spore traps for the current study.

The rotating-arm spore traps were constructed within the context of our study based on an adaptation of the manual by Quesada et al. (2018). The air sampling rate per minute (V in L/min) for one arm of the rotating-arm spore trap is calculated using the equation formulated by McCartney et al. (1997): $V = A \times\ 2R\ \times\ \pi\ \times\ rpm$, with A= rod area ($m^2$), R= radius of the arm (m), rpm= rotations per minute. The radius of our constructed rotating-arm spore trap was 25 cm, the rotations per minute were 30 (given by the motor), and the sampling area was 25 x 40 mm. Since the air sampling rate of our rotating-arm spore trap is relatively low, we decided to run the traps continuously for periods of at least ten days.

The rotating-arm spore trap consisted of two rods connected to a battery-powered motor (spinart$^{TM}$ Lightweight Hanging Motor, 30 rpm, operated with a LR20 1.5 V battery). Each rod held a microscope slide on which a Vaseline coated film strip was mounted and fixated with 25 mm foldback clips (Maul, Bad König, DE). The microscope slide was used to provide further rigidity to the spore trapping surface, the film strip, which had to be flexible enough to be folded into a 2 mL tube for DNA extraction. The rods were made of 15 cm long segments of galvanized steel wire with 1.1 mm in diameter. The rods were mounted on the rotating part of the battery motor using ring terminal cable lugs (0.15-1.5 x 5.4 mm). A 30 x 30 cm shield made of aluminum coated with polyfilm (Universal laser-film, Fiber Optic & $CO_2$ Technologies) was used as a rain protection. The shield was secured to the bottom of the battery motor using two screws. For this, a hole with 85 mm in diameter was drilled into the middle of the shield for letting the rotating element through. A 40 mm diameter hole was drilled 15 mm distance from the middle on both sides for the two screws. Two matching holes were drilled into the bottom part of the battery motor by unscrewing the motor first and making sure a nut would fit inside the motor without touching any rotary knob. The rods were made by cutting a galvanized steel wire of 1.1 mm diameter into 15 cm segments. The rods were mounted on the rotating part of the battery motor using a ring terminal cable lug (0.15-1.5 mm x 5.4 mm). The cable lugs were held in place by nuts (M 4, Inox A2) mounted on the rotating end of the motor into which a thread was cut.

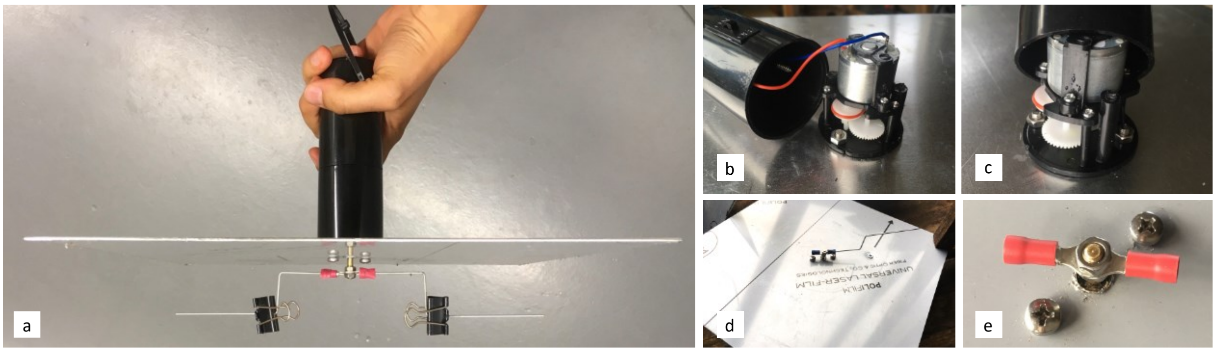

**Construction of the rotating-arm spore trap.** a) Finished rotating-arm spore trap, b)-c) motor of the rotating-arm spore traps screwed to an aluminum shield, d) underside of the rotating-arm spore trap (one arm fixed with a ring terminal lug to the rotating part of the motor), e) closer view of the ring terminal lugs.

**Supplementary Materials and Methods S3.** Preparation of the vaseline coated film as the spore trapping surface.

For the Mycotrap, an A4 (21 x 29.7 cm) transparent film sheet, as used for laser printers, was cut lengthwise into 2.5 x 29.7 cm film strips. The film strips were fixed with tape on a black plastic tray. Each film strip was evenly sprayed with Vaseline dissolved into pentane (Vaseline:pentane, 1:2) (Lacey and West 2006) using a spraying gun (DeVilbiss GPi multi-purpose spray gun, Devilbiss UK) and let to dry. The final amount of Vaseline on the film was approximately 0.01 g Vaseline per $cm^2$. The Vaseline coated film strip was mounted on the drum of the Mycotrap together with a paper strip onto which a time scale was printed. The beginning of the Vaseline coated film strip was aligned with the starting time of the sampling as well as the indent on the cover of the drum indicating the location of the sampling opening.

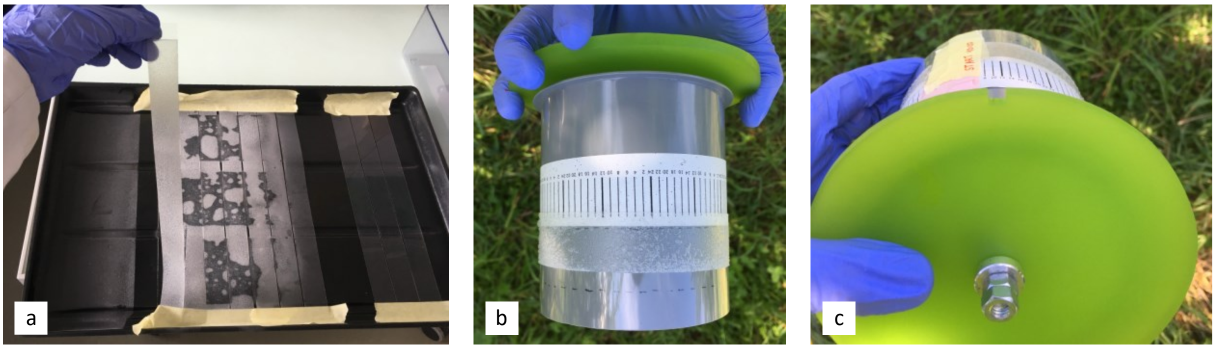

**Preparation of spore trapping surface for the Mycotrap.** a) Spraying dissolved Vaseline onto transparent film strips, b) mounting the Vaseline coated film strip onto the cylinder of the Mycotrap underneath a paper strip with the time scale, c) placing the orifice at the start of the Vaseline strip.

For the rotating-arm spore traps, an A4 transparent film sheet was cut into 2.5 x 21 cm strips. Only the middle section of these strips was sprayed with Vaseline as described above. Each strip was cut into two halves across its width. Each half was cut into a 76 mm (length of a microscope slide) long strip to obtain a segment, which on one end was coated with Vaseline covering at least 40 mm while the other end was without Vaseline in order to tape it on a clean microscope slide.

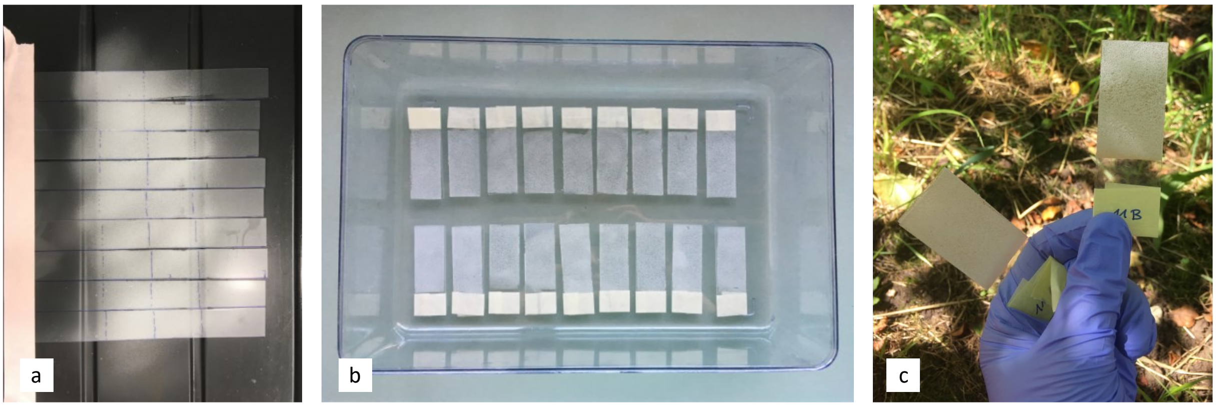

**Preparation of spore trapping surface for the rotating arm traps.** a) Vaseline coated plastic strips sprayed with Vaseline. b, c) Strips were fixed with tape onto microscope glass slides.

**Supplementary Materials and Methods S4.** Protocol for DNA extraction with CTAB.

The following protocol is a modification of the CTAB (Cetyltrimethylammonium-bromide) extraction as described in Angelini et al. (2001).

**Buffers**

| **CTAB Extraction Buffer (HCl pH 8.0)** | | **100 mL** |
|---|---|---|
| Tris (Roth 5429.3) | 100 mM | 1.21 g |
| NaCl (Roth 3957.1) | 1.4 M | 8.19 g |
| $Na_2EDTA$ (Roth 8043.1) | 50 mM | 1.86 g |
| CTAB (Roth 9161.2) | 2% | 2.0 g |
| PVP K25 (Roth 4606.2) | 1% | 1.0 g |
| HCl (5 N) pH 8.0, | | |
| Buffer stable for 2 years at room temperature. | | |
| β-Mercaptoethanol (added just before use) | 0.2% | 0.2 mL |

| **TE (HCl pH 8.0) 10x** | | **100 mL** |
|---|---|---|
| Tris (Roth 5429.3) | 100 mM | 1.21 g |
| $Na_2EDTA$ (Roth 8043.1) | 10 mM | 0.37 g |

**Procedure**

1. Put 0.5 to 0.6 g leaf sample in one of the compartments of an extraction bag (Bioreba AG, Reinach, Switzerland). Add β-Mercaptoethanol freshly to the necessary amount of extraction buffer, and add 3 to 5 mL of it to each sample. Homogenize with the HOMEX (Bioreba AG, Reinach, Switzerland). (When using frozen samples, make sure that they are processed immediately without thawing before adding the extraction buffer).

2. Transfer 1 mL of the extract into a 2 mL tube with a screw cap and incubate at 70°C for 20 min.

3. After cooling, add 0.8 mL Chloroform (Roti Cl) in the hood and vortex for 3x 2 sec at full speed by holding the tube at the top. This allows the liquid level reach to the top of the tube and ascertains a thorough mixing of the two phases.

4. Centrifuge at 20,000 x g for 2 min.

5. Add 0.6 mL Isopropanol into a new 1.5 mL reaction tube and then transfer 0.8 mL of the clear supernatant of each sample. Avoid lower and interphase. Mix well by inverting the tubes upside down 4 to 5 times (Do not vortex!). Incubate 1 h at room temperature or overnight at -20°C to let the DNA precipitate.

6. Centrifuge at 20,000 x g for 2 min, discard the supernatant and wash the DNA pellet once with 0.8 mL EtOH 70%. Centrifuge again at 20,000 x g for 1 min and discard the supernatant. Be careful not to lose the pellet in this latter step. It is recommended to pour off the supernatant (70% EtOH) within maximal 60 s after centrifugation.

7. Evaporate the EtOH in the pellet by placing the open tubes for 5 to 15 min in the heat block at 70°C.

8. Dissolve the DNA pellet in 100 µL of TE 0.1x by flicking the tube gently with a finger. Do not vortex – at least not excessively. Assessment of DNA concentration and purity by Nanodrop. Keep the DNA extracts at 4°C for use within a couple of weeks or at -20°C for long-term storage.

**Supplementary Materials and Methods S5.** APA9 plasmid as reference for DNA extraction efficiency.

The plasmid APA9 (vector pUC19 with African cassava mosaic virus insert; GenBank accession number AJ427910) was multiplied in *Escherichia coli*. *E. coli* from the -80°C long-term storage was grown on LB plates with chloramphenicol and tetracycline. The plasmid was extracted from an overnight culture in liquid LB medium (14 h, 37°C, 120 rpm on an Ecotron shaker, Infors HT®, Basel, CH) with the peqGOLD Plasmid Miniprep Kit II (peqlab®, Erlangen, DE) according to the manufacturer's instruction. Next, the plasmids were linearized by adding 1 µL of the restriction enzyme NotI (R0189S, 10,000 U/mL, New England BioLabs®, MA, USA), 5 µL of NEBuffer 3.1 (B7203S, 10X Concentration, New England BioLabs®, MA, USA), and 34 µL qPCR grade water to 10 µl APA9 DNA ($10^{10}$ copies APA9). The DNA concentration was measured with Qubit® (Thermo Fisher Scientific, MA, USA) and copy numbers calculated using the Endmemo copy number calculator (http://endmemo.com/bio/dnacopynum.php). Aliquots of $10^6$ copies APA9/µL were stored at -20°C until usage.

All spore trap samples were spiked with $10^7$ copies of linearized APA9 (10 µL of a $10^6$ copies APA9/µL solution). The qPCR for the plasmid was multiplexed with the qPCR for *D. coronariae* using a TaqMan based qPCR. The $C_q$ values were normalized using the formula published by Von Felten et al. (2010):

$$\text{Normalized } C_q \text{ value of sample X} = \frac{C_q \text{value of sample X}}{C_q \text{value of internal standard in sample X}} \times \text{average } C_q \text{ value of internal standard in all samples}$$

**Supplementary Materials and Methods S6.** Installation of a Mycotrap on a leaf litter deposit.

In 2019, a Mycotrap was placed in the field on a mesh cage depot filled with heavily *Dc* infested leaf litter of the variety 'Remo' at the Competence Center for Fruit Crops at the Lake of Constance (KOB).

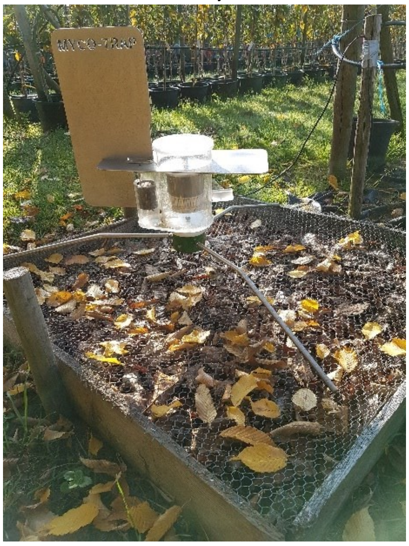

**Spore trap above leaf litter deposit at KOB.**

### Supplementary Tables

**Supplementary Table S1.** Fungal strains used in this study.

| Strain name | Other names | Origin | Reference |
|---|---|---|---|
| *Diplocarpon coronariae* CH01 | *Marssonina coronaria, Diplocarpon mali* | Apple, Switzerland | (Oberhänsli et al. 2021) |
| *Diplocarpon coronariae* CH02 | *Marssonina coronaria, Diplocarpon mali* | Apple, Switzerland | (Oberhänsli et al. 2021) |
| *Diplocarpon coronariae* CH03 | *Marssonina coronaria, Diplocarpon mali* | Apple, Switzerland | (Oberhänsli et al. 2021) |
| *Diplocarpon coronariae* Japan NBRC 30405 | *Marssonina coronaria, Diplocarpon mali* | Apple, Japan | NBRC 30405, NITE, Tokyo, Japan |
| *Diplocarpon coronariae* Korea GW3 | *Marssonina coronaria, Diplocarpon mali* | Apple, Korea | (Oberhänsli et al. 2021) |
| *Diplocarpon rosae* CBS 163.31 | *Marssonina rosae* | *Rosa,* country unknown | Westerdijk Fungal Biodiversity Institute (Utrecht, The Netherlands) |
| *Diplocarpon mespili* CBS 166.28 | *Entomosporium mespili* | Cydonia leaf, England | Westerdijk Fungal Biodiversity Institute (Utrecht, The Netherlands) |
| *Diplocarpon earlianum* CBS 162.32 | *Marssonina fragariae* | *Fragaria,* country unknown | Westerdijk Fungal Biodiversity Institute (Utrecht, The Netherlands) |
| *Drepanopeziza tremulae* MUCL 8333 | *Marssonina tremulae, Marssonina brunnea* | Poplar leaf, Switzerland | Belgium co-ordinated collection of micro-organisms (BCCM/MUCL) |
| *Gnomonia leptostyla* DSMZ 1462 | *Ophiognomonia leptostyla, Marssonina juglandis* | Walnut leaves, France | DSMZ-German Collection of Microorganisms and Cell Cultures GmbH |
| *Neofabraea alba* MUCL 46541 | *Pezicula alba, Phlyctema vagabunda* | No information | Belgium co-ordinated collection of micro-organisms (BCCM/MUCL) |
| *Neofabraea malicorticis* MUCL 31836 | *Pezicula malicorticis* | No information | Belgium co-ordinated collection of micro-organisms (BCCM/MUCL) |
| *Neonectria galligena* MUCL 6129 | *Cylindrocarpon mali* | Fruit, Belgium | Belgium co-ordinated collection of micro-organisms (BCCM/MUCL) |
| *Monilinia fructigena* MUCL 18345 | *Monilia fructigena* | Fruit, Belgium | Belgium co-ordinated collection of micro- |

| | | | |
|---|---|---|---|
| | | | organisms (BCCM/MUCL) |
| *Penicillium expansum* MUCL 20453 | | Apple fruit, Belgium | Belgium co-ordinated collection of micro-organisms (BCCM/MUCL) |
| *Penicillium digitatum* MUCL 46389 | | Orange, Belgium | Belgium co-ordinated collection of micro-organisms (BCCM/MUCL) |
| *Colletotrichum gloesporioides* 2020-20 | *Glomerella cingulata* | Apple fruit, Switzerland | This study |
| *Botrytis cinerea* TwA86 | *Botryotinia fuckeliana* | Apple fruit, Switzerland | This study |
| *Fusarium oxysporum* Ab1c56 | | Apple fruit, Switzerland | This study |
| *Phomopsis* sp. Arc77 | | Apple fruit, Switzerland | This study |
| *Venturia inaequalis* | *Spilocaea pomi* | Spores collected from apple leaves, Switzerland | This study |

**Supplementary Table S2.** Spore traps and sampling periods for monitoring the temporal and spatial spore dispersal in an apple orchard in 2019[a].

| Sampling period (date)[b] | Spore traps | | | Bait plants[e] |
|---|---|---|---|---|
| | Mycotrap 1[c] | Mycotrap 2[d] | Rotating-arm spore trap | |
| 09.05.-14. 05. | x | | | x |
| 14. 05.-17.05. | x | | | |
| 17.05.-22.05. | x | | 2 prototypes | x |
| 22.05.-28.05 | x | | x | x |
| 28.05.-03.06. | x | | x | x |
| 03.06-07.06 | x | x | x | x |
| 07.06-13.06 | x | x | x | x |
| 13.06-19.06 | x | x | x | x |
| 19.06-25.06 | x | x | x | |
| 25.06.-01.07. | x | x | x | |
| 01.07-05.07. | x | x | x | |
| 05.07-10.07. | x | | | |
| 10.07.-17.07. | x | | | |
| 17.07.-30.07. | x | | | |

[a] The field trial lasted from 9 May until 30 July 2019. x, trap was running in this sampling period.
[b] The traps were run permanently, but at the end of each sampling period, the Vaseline coated strips had to be exchanged in case of the Mycotraps. The Vaseline coated strips in the rotating-arm spore traps were only exchanged every other sampling, i.e. every 10 to 12 days.
[c] Mycotrap 1 was installed on the ground.
[d] Mycotrap 2 was installed in the tree crown.
[e] Each set of bait plants consisted of three 'Topaz', three 'Gala' and three 'Kiku' potted apple trees. After each sampling period, the bait plants were exchanged. Bait plants coming from the field were incubated in a foliar tunnel and monitored for apple blotch symptom development.

**Supplementary Table S3**. Overview of spore traps and sampling periods for monitoring the temporal and spatial spore dispersal in an apple orchard in 2020 [a].

| Sampling period (date)[b] | Spore traps | | | |
|---|---|---|---|---|
| | Mycotrap 1[c] | Mycotrap 2[d] | Rotating-arm spore trap | Bait plants[e] |
| 28.2.-13.3. | x | x | x | |
| 13.3.-27.3. | x | x | x | x |
| 27.3.-10.4. | x | x | x | x |
| 10.4.-24.4. | x | x | x | x |
| 24.4.-8.5. | x | x | x | x |
| 8.5.-22.5. | x | x | x | x |
| 22.5.-5.6. | x | x | x | x |
| 5.6.-19.6. | x | x | x | x |
| 19.6.-3.7. | x | x | x | x |
| 3.7.-17.7. | x | x | x | x |
| 17.7.-30.7. | x | x | x | x |

[a] The field trial lasted from 28 February until 30 July 2020. In the first sampling period, no bait plants were placed in the orchard since neither bait plant nor trees in the orchard had leaves at that time. x, trap was running in this sampling period.
[b] The traps were ran permanently, but at the end of each sampling period, the Vaseline coated strips were exchanged in the Mycotraps. The Vaseline coated strips in the rotating-arm spore traps were only exchanged every other sampling period, i.e. always after 14 days.
[c] Mycotrap 1 was installed on the ground.
[d] Mycotrap 2 was installed in the tree crown.
[e] Each set of bait plants consisted of five 'Topaz' trees. After each sampling period, the bait plants were exchanged. Bait plants from the field were incubated in the greenhouse and monitored for apple blotch symptom development.

**Supplementary Table S4.** Primer and probe Dc_09 specifically amplify all *Diplocarpon coronariae* isolates.[a]

| **Sample** | **Cq value[b]** | | | |
|---|---|---|---|---|
| | **Dc_09** | | **ITS1F/2R** | |
| *D. coronariae* CH01 | 25.0 | ± 0.2 | 27.0 | ± 0.7 |
| *D. coronariae* CH02 | 24.2 | ± 0.1 | 25.0 | ± 0.4 |
| *D. coronariae* CH03 | 22.3 | ± 0.1 | 22.8 | ± 0.3 |
| *D. coronariae* Japan NBRC 30405 | 21.9 | ± 0.7 | 24.4 | ± 0.6 |
| *D. coronariae* Korea GW3 | 26.9 | ± 0.2 | 25.4 | ± 0.2 |
| *Diplocarpon mespili* CBS 166.28 | ND | | 16.2 | ± 0.2 |
| *Diplocarpon rosae* CBS 163.31 | ND | | 12.1 | ± 0.2 |
| *Diplocarpon earlianum* CBS 162.32 | ND | | 15.3 | ± 0.1 |
| *Drepanopeziza tremulae* MUCL 8333 | ND | | 14.9 | ± 0.4 |
| *Gnomonia leptostyla* DSMZ 1462 | ND | | 16.4 | ± 0.1 |
| *Neofabraea alba* MUCL 46541 | ND | | 14.9 | ± 0.3 |
| *Neofabraea malicorticis* MUCL 31836 | ND | | 14.8 | ± 0.3 |
| *Neonectria galligena* MUCL 6129 | ND | | 15.9 | ± 0.1 |
| *Monilia fructigena* MUCL 18345 | ND | | 19.1 | ± 0.1 |
| *Penicillium expansum* MUCL 20453 | ND | | 11.2 | ± 0.1 |
| *Penicillium digitatum* MUCL 46389 | ND | | 10.3 | ± 0.1 |
| *Colletotrichum gloeosporioides* 2020-20 | ND | | 17.0 | ± 0.1 |
| *Botrytis cinerea* TwA86 | ND | | 18.1 | ± 0.3 |
| *Fusarium oxysporum* Ab1c56 | ND | | 14.9 | ± 0.2 |
| *Phomopsis* sp. Arc77 | ND | | 14.6 | ± 0.1 |
| *Venturia inaequalis* | ND | | 26.8 | ± 0.1 |
| healthy apple leaf | ND | | 23.1 | ± 0.1 |
| No template control | ND | | ND | |

[a] DNA was amplified by a *D. coronariae* specific TaqMan qPCR using primers and probe Dc_09 and by a SYBR green based qPCR using the general fungal primers ITS1F/ITS2R, which served as control.

[b] Cq = quantification cycle. Mean ± sdev of three technical replicates are shown. ND, no amplicon detected.

**Supplementary Table S5.** Detection of *Diplocarpon coronariae* DNA on fruit mummies in winter.[a]

| Sample name | Apple cultivar[b] | Norm $C_q$ TaqMan[c] | *D. coronariae* |
|---|---|---|---|
| Fruit 3-15 | Otava | ND | negative |
| Fruit 15-15 | Topaz | 37.26 | positive |
| Fruit S4 | Topaz | 40.12 | positive |
| Fruit 8-18 | Florina | 43.06 | negative |
| Fruit S2 | Florina | 33.65 | positive |
| Fruit 10-18 | Schneider | ND | negative |
| Fruit 9-16 | Schneider | 40.38 | positive |
| Fruit S7 | Blauacher | ND | negative |
| Fruit S3 | Blauacher | ND | negative |
| Fruit S7 | Blauacher | ND | negative |
| Fruit S5 | Rubinola | ND | negative |
| Fruit S6 | Rubinola | ND | negative |
| Fruit S10 | Rubinola | ND | negative |
| Leaf 8-13 | Florina | 25.83 | positive |

[a] Fruit samples were collected from apple trees in the Rickenbach orchard (Switzerland) on 28 February and 13 March 2020.
[b] 'Topaz', 'Otava', and 'Florina' trees were heavily infected with *D. coronariae* in 2019, while 'Blauacher' and 'Schneider' trees were less affected by apple blotch, and 'Rubinola' trees showed no symptoms.
[c] Cq (quantification cycle) values result from a TaqMan qPCR with primers and probe Dc_09. APA9 plasmid was used to normalize for DNA extraction efficiency. ND, not detected.

**Supplementary Table S6.** Kendall rank correlation test between daily spore number and precipitation for May, June, and July 2019 and 2020.

| Year | Month | rank correlation coefficient | p-value | significance |
|---|---|---|---|---|
| 2019 | May | -0.32 | 0.045 | significant |
| | June | -0.03 | 0.850 | not significant |
| | July | 0.15 | 0.319 | not significant |
| 2020 | May | -0.03 | 0.867 | not significant |
| | June | 0.28 | 0.056 | not significant |
| | July | -0.10 | 0.492 | not significant |

Supplementary Figures

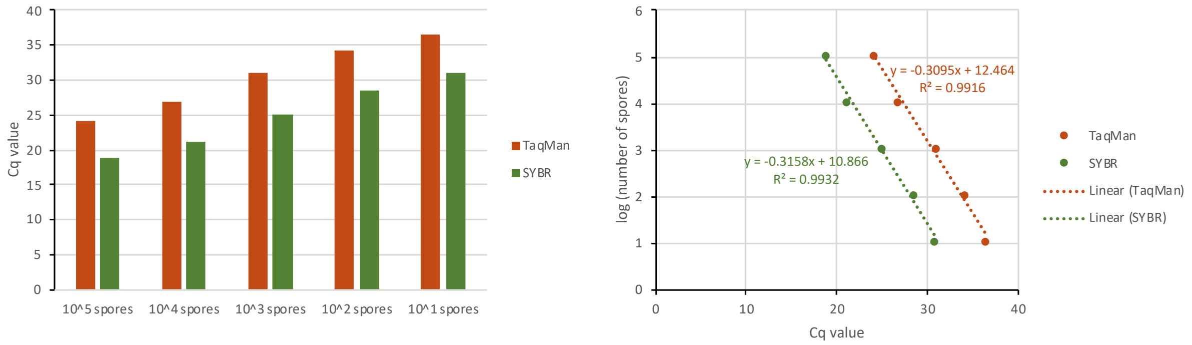

**Supplementary Fig. S1.** Comparable sensitivity with SYBR based qPCR and TaqMan based qPCR. The standard curve with $10^5$, $10^4$, $10^3$, $10^2$ and $10^1$ *D. coronariae* conidia extracted from Vaseline coated film tested with SYBR Green I qPCR (E = 100%) and TaqMan qPCR (E = 105%) (detection limit for SYBR $C_q$ = 35, for TaqMan $C_q$ = 40).

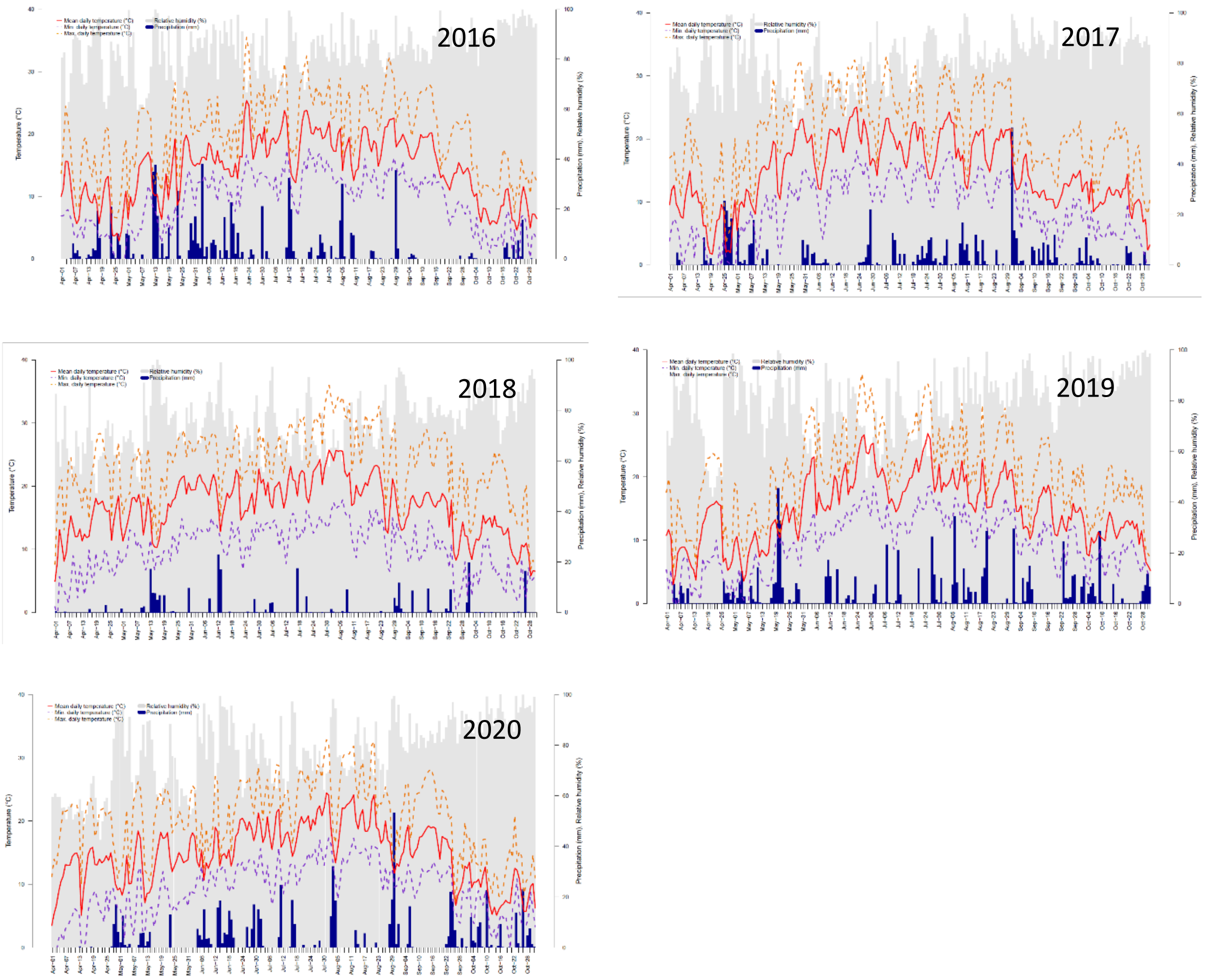

**Supplementary Fig. S2.** Weather data for the location KOB and the year 2016 to 2020. The weather data includes temperature (°C), relative humidity (%), and precipitation (mm). Weather data from the weather station in Bavendorf (47°76'84.4"N 9°55'99.0"E), Germany, close to the Competence center for fruit crops at the Lake of Constance (Kompetenzzentrum Obstbau Bodensee, KOB), located in Ravensburg-Bavendorf, Germany.

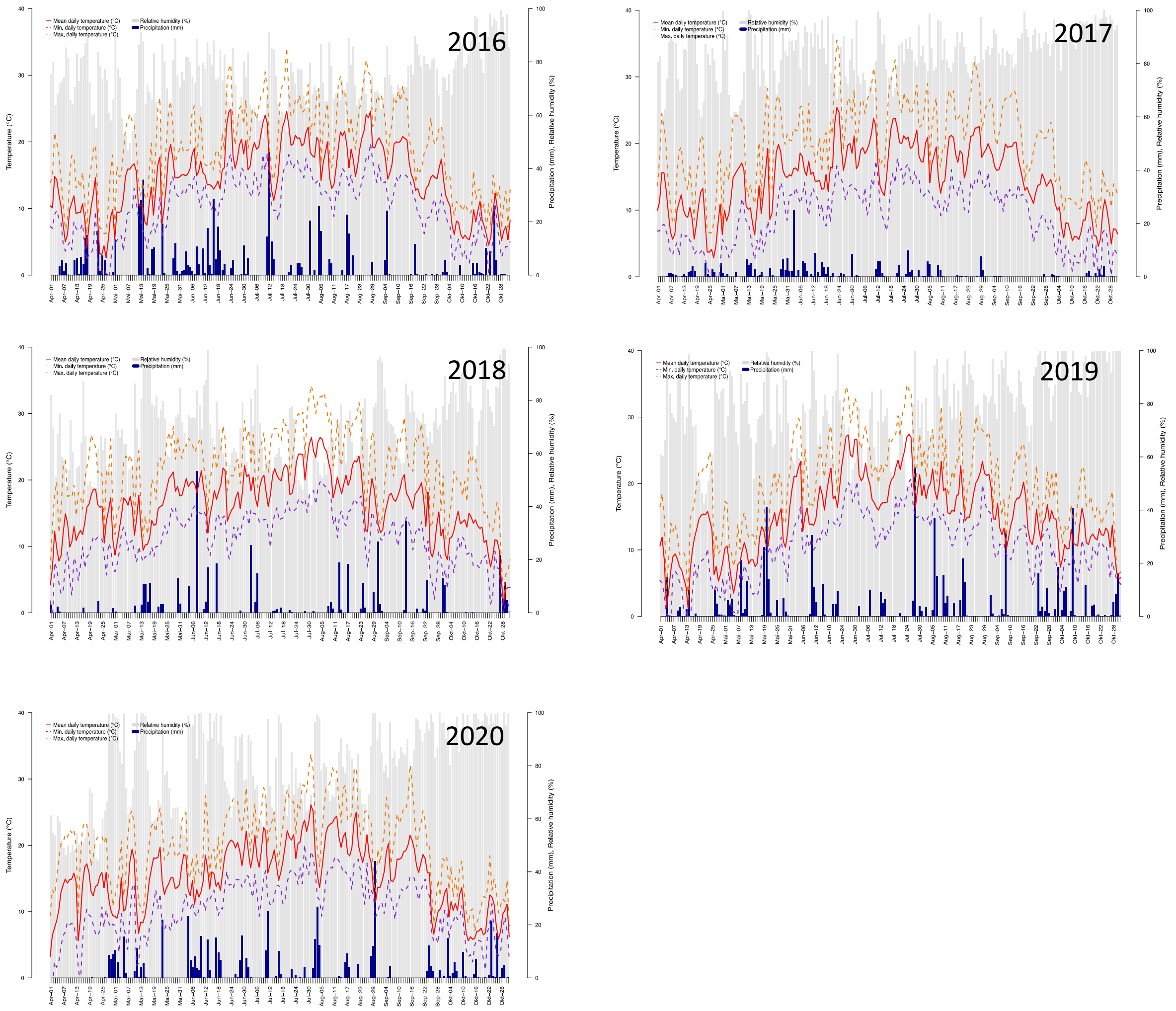

**Supplementary Fig. S3.** Weather data for the location Rickenbach and the years 2016 to 2020. The weather data includes temperature (°C), relative humidity (%), and precipitation (mm) (data from Agrometeo, location “Liebensberg”, Zurich (47°53'38.1"N 8°83'73.8"E)).

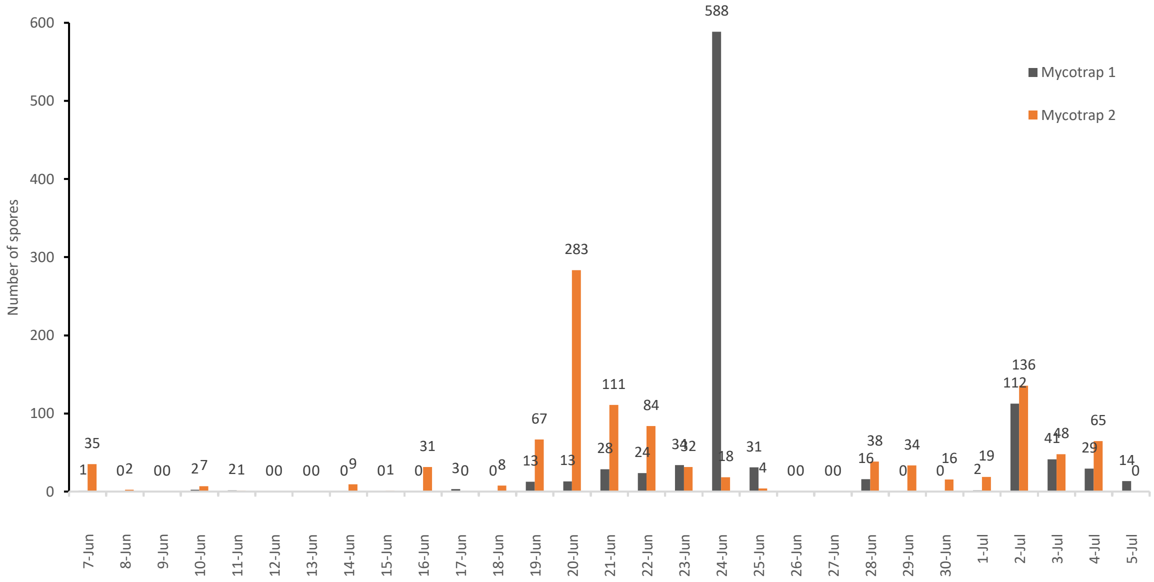

**Supplementary Fig. S4.** Comparison of spores caught by a Mycotrap placed on the ground vs. in the tree crown (2019). From 7 June to 5 July, a second Mycotrap (Mycotrap 2) was placed inside the tree crown of an 'Otava' apple tree next to the first Mycotrap (Mycotrap 1), which was placed on the ground (opening orifice at 20 cm above ground).

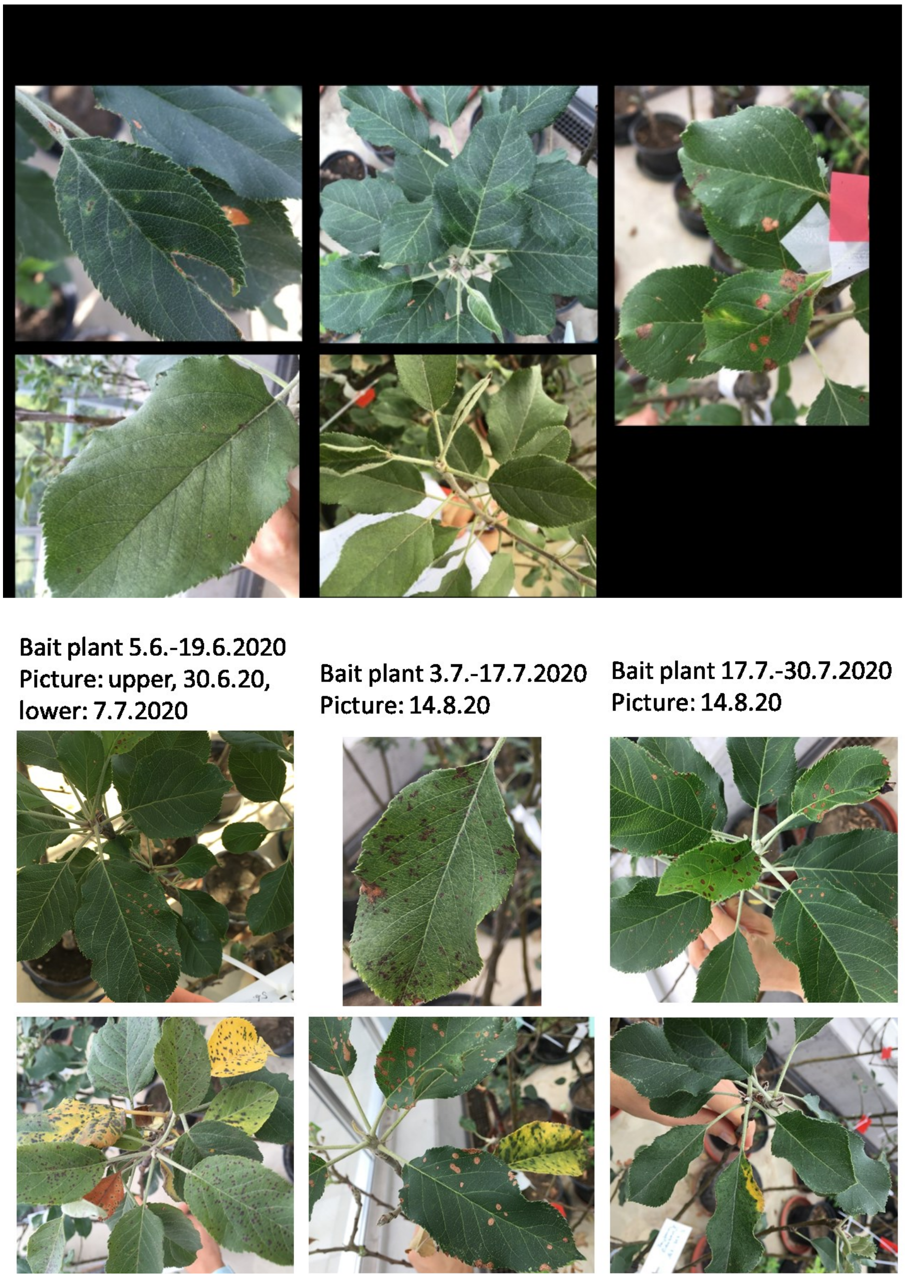

**Supplementary Fig. S5.** Apple blotch symptoms on bait plants (potted 'Topaz' apple trees) in 2020. During the field experiment five two-year-old trees were placed close to the Mycotraps in the apple orchard. These bait plants were exchanged every two weeks, transferred to the greenhouse and symptom development was assessed. First bait plants developed symptoms, from 24 April until 8 May, but symptoms were rather weak. The next series (8 May until 22 May) did not develop symptoms. After 22 May, all bait plants were infected with *D. coronariae* and exhibited strong symptoms. Pictures of a few representative trees are shown. Leaves with ambiguous symptoms were tested by qPCR.

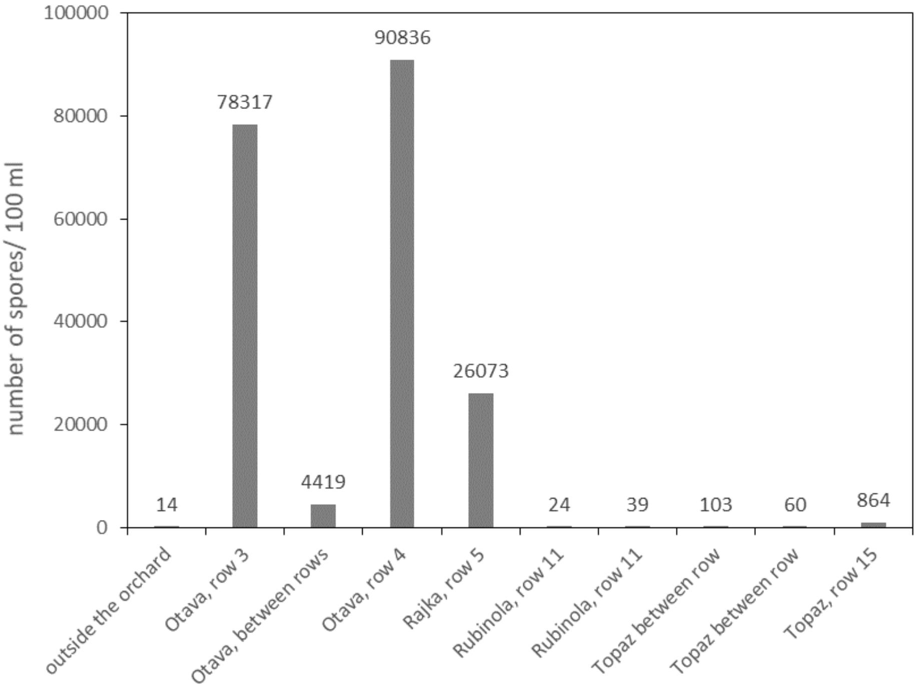

**Supplementary Fig. S6.** Large amounts of *D. coronariae* conidia in rainwater below highly infected trees. Rainwater was collected below or between tree rows from 24 to 25 September 2020. The water was filtered through a cellulose acetate filter. The DNA was extracted from the filter, and *D. coronaria*e spores were quantified by qPCR. Spore numbers were calculated based on a standard curve with known amounts of *Dc* conidia.

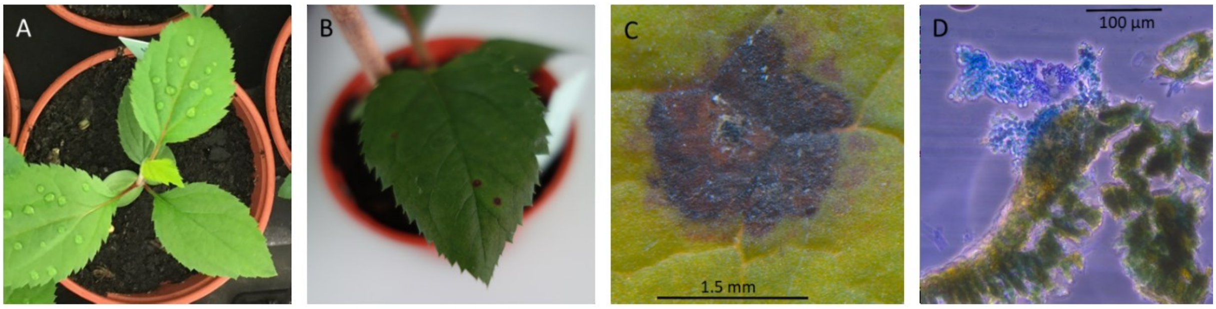

**Supplementary Fig. S7**. Apple seedling infected with *D. coronariae* (*Dc*) from overwintered fruit. *Dc* conidia were obtained from an infected 'Otava' fruit collected in the Rickenbach orchard in September 2020 and stored in the fridge until March 2021. A conidia suspension was pipetted onto the apple leaf (A), and symptoms were photographed after one month (B). Acervulus formation was photographed after three months under a stereomicroscope (C) Stereomicroscopic view of an acervulus that formed on the leaf.
